# Lateral flow cell-free transcriptional assay for contaminant detection

**DOI:** 10.1101/2025.08.20.670534

**Authors:** Matias Villarruel Dujovne, Carolina Silva, Fatima Alvarez Rocco, Gustavo Ghiglieri, Mariana Riesgo, Daiana A. Capdevila, Ana Sol Peinetti

## Abstract

Cell free transcriptional biosensors are emerging as a powerful technology, offering enhanced capabilities for detecting chemical contaminants in settings where traditional analytical techniques fall short of societal needs. However, the sensitivity of these biosensors for many critical chemical contaminants often remains insufficient to meet regulatory detection limits, while maintaining portability and ensuring selectivity. In this work, an *in vitro* transcription (IVT) based lateral flow assay (LFA) is reported, which is 1000 times more sensitive than reported IVT portable detection methods. Through combining the amplification power of IVT with nucleic acid LFA-based avidity effects, we develop rapid, ultra-sensitive turn-on sensors. Our sensors meet regulatory detection limits for lead, a particularly challenging metal ion contaminant, in untreated spring water samples collected from a basin that exemplifies the environmental monitoring challenges. Additionally, we show that by optimizing the IVT conditions, we can tune the limit of detection to produce an on-off signal reaching the WHO regulation threshold and others. Finally, we evaluate the selectivity of different metal-responsive repressors in the context of *in vitro* biosensing and discuss how to identify potentially hazardous samples with an easy-to-use and point-of-care assay.

## 1. INTRODUCTION

Cell-free transcriptional biosensors are emerging as low-cost, user-friendly, and portable tools with significant potential for medical diagnostics and environmental monitoring (Slomovic et al., 2015; Jung et al., 2020; Sadat Mousavi et al., 2020; Jung et al., 2022; Phillips et al., 2023). Successful *in vitro* transcription systems have been engineered to respond to a broad range of analytes (from metal ions to small molecules and proteins) which elicit a transcriptional response from allosteric transcription factors (aTFs)(Capdevila et al., 2024; Ekas et al., 2024a; Jung et al., 2020; Wang et al., 2025) and nucleic acid regulatory elements (Bracaglia et al., 2023; Chou and Shih, 2019; Lee et al., 2024; Miceli et al., 2025; Patino Diaz et al., 2022; Thavarajah et al., 2020). In particular, the RNA Output Sensors Activated by Ligand Induction (ROSALIND) platform employs apo-repressors as transcription factors from different structural families to sense a wide array of metal and organic contaminants in water (Jung et al., 2020). The platform combines a linear DNA template with a bacteriophage T7 RNA polymerase (RNAP) promoter and a DNA operator sequence that binds the transcription factor of choice together with the purified polymerase and aTF. These elements regulate the transcription of an RNA aptamer that, when bound to an organic dye, produces a fluorescent signal. Furthermore, the fluorescent output can be detected using portable handheld illuminators created with 3D printed parts, simple electronic components and inexpensive LEDs enabling on-site detection of these metal ions and small molecules (Jung et al., 2020). However, while promising, there are still several challenges that remain unresolved in terms of the limit of detection, selectivity, and portability for many chemical contaminants. For example, existing lead sensors still struggle to achieve the low limits of detection required for regulatory compliance (Ekas et al., 2024a; Jung et al., 2020; Li et al., 2025). Also, it is not clear whether the appropriate selectivity in complex real samples can be achieved by lead sensitive aTFs, as most transcription factors may only be specific for particular chemical species in the cellular context and are expected to be more promiscuous *in vitro* (Clough et al., 2025). Addressing these limitations is critical for achieving the full potential of cell-free biosensors in practical applications (see Supplementary text 1).

In an *in vitro* transcription (IVT) based biosensor, the limit of detection of a particular chemical contaminant is largely determined by the concentration of the repressor necessary to achieve full repression in a ligand-free buffer. If the concentration of the repressor required to avoid background signal is too high, the sensor often fails to meet practical or regulatory requirements for the contaminant the repressor is responsive to. To overcome this limitation different strategies have been reported. On one hand, to decrease the amount of DNA template required and still have a detectable output signal, IVT products have been amplified through an isothermal signal amplification circuit called polymerase strand recycling (PSR) that leverages T7 RNA polymerase off-target transcription to recycle nucleic acid inputs within DNA strand displacement circuits (Li et al., 2025). However, amplifying the IVT reaction product by isothermal amplifications can generate higher background signal and the need of modified DNA oligos may increase the overall reaction cost. On the other hand, it is possible to tune the DNA binding affinities of the aTFs to improve sensitivity. This was achieved by a high-throughput platform that allows for the synthesis and discovery of mutant aTFs with improved sensitivity (Ekas et al., 2024a, 2024b). However, this strategy also led to increased background signal, which may be problematic when it comes to implementing these proteins in point-of-care sensors. Moreover, most examples of point-of-care sensors involve fluorescence detection. While this often allows for better sensitivity than colorimetric readouts, it also has disadvantages due to quenching effects from interferences in complex samples and the difficulties obtaining affordable and sensitive portable fluorescence devices (Dias et al., 2025; Quero et al., 2024). Therefore, the IVT based sensors would benefit from amplification strategies that involve signal transduction methods leading to a robust signal even at low concentrations of template and repressor, while maintaining the advantage of rapid, instrument-free, on-site detection.

One of the most successful biosensors available to the public are lateral flow assays, with lateral flow immunoassays commercially introduced over 30 years ago, enabling the detection of a wide variety of antigens (Bahadır and Sezgintürk, 2016; Peri Ibáñez et al., 2024; Parolo et al., 2020). LFA methods for selectively detecting metal ion contamination have been developed using antibodies (López_Marzo et al., 2013), aptamers (Srinivasan et al., 2023; Wu et al., 2017) and DNAzymes (Mazumdar et al., 2010), however the limit of detection of many developments do not meet regulatory needs. In recent years, lateral flow assays have also been adapted for nucleic acid detection, opening the possibility of coupling nucleic acid amplification with LFA detection (Phillips et al., 2023; Wang et al., 2020). Several approaches have been employed, primarily focusing on two main strategies. The first generally involves amplifying nucleic acids using PCR or other amplification techniques, with different hapten-tagged primers. In this technique, called Nucleic Acid Lateral Flow Immunoassay (NALFIA), the hapten-labeled DNAs or other tagged nucleotides are recognized by antibodies or streptavidin on the test and control lines (Fong et al., 2000; Jauset-Rubio et al., 2016; Kellner et al., 2019; Lee et al., 2025). This approach has been incorporated in CRISPR-Cas based sensors interfaced with LFA transduction methods, generating portable pathogen nucleic acids detection (Myhrvold et al., 2018; Patchsung et al., 2020; Tang et al., 2022; Wang et al., 2020). The second approach, called Nucleic Acid Lateral Flow (NALF) is based only on nucleic acid hybridization as the recognition event. This is accomplished by using two types of oligonucleotide probes (capture probes and reporter probes) to detect single-stranded sequences, where both probes must directly hybridize with the target sequence in the test line to obtain a colorimetric signal (Roskos et al., 2013). Despite the advantages of multiplexing in the same cassette test, there are fewer examples of applications of this approach. This is possibly due to the hybridization kinetics in lateral flow being more complex compared with the formation of hapten-antibody complexes (Jauset-Rubio et al., 2016). As functional DNA engineering strategies continue to develop, it is likely that NALF strategies that directly detect DNA will become more frequently used than NALFIA strategies that require additional antibodies and labelled DNA. We aim here to fill a gap in the development of NALF based tests by introducing avidity-based probes that harness optimized kinetics enabling enhanced detection.

Beyond limit of detection and easy to use readouts, the selectivity of cell-free transcriptional biosensors is directly determined by the specificity of the aTFs in the absence of any cellular context, which does not necessarily reach the analytical performance required for water monitoring (Capdevila et al., 2024; Jung et al., 2020). While in cellular contexts many aTFs show a remarkable selectivity towards different chemical species and particularly transition metal ions (Capdevila et al., 2024, 2017; Clough et al., 2025; Foster et al., 2022; Lenner et al., 2025; Osman et al., 2019), their behavior under *in vitro* conditions with varying metal ion concentrations has not been thoroughly investigated. The observed degrees of cross-reactivity cast doubts on the conclusions drawn from initial studies (Beabout et al., 2021; Jordan et al., 2020; Jung et al., 2020). For example, CadC repressors have shown cross-reactivity between lead and zinc *in vitro* (Jung et al., 2020), which is also compatible with the biochemical characterization of metal and DNA binding (Busenlehner et al., 2002, 2001). Thus, to further develop the use of aTFs for IVT based sensors, it is critical to evaluate the level of crosstalk between different chemical species *in vitro*. This characterization is usually limited to obtaining metal binding (Busenlehner et al., 2002, 2001; Tóth et al., 2024) or metal preference (Clough et al., 2025) parameters. Recently, ROSALIND was employed as a tool to determine the residues involved in metal coordination (Kim et al., 2024). Here we use ROSALIND as a tool to define to what extent the *in vitro* selectivity is sufficient to distinguish divalent cations from the Irving-Williams series (Irving and Williams, 1953), also including xenobiotics such as lead.

In this work, we designed nucleic acids to detect IVT products using an avidity-based NALF assay and developed a portable detection method for metal ion pollutants. Our approach combines the amplification power of IVT with avidity effects derived from nucleic acid LFA to achieve rapid and ultra-sensitive turn-on sensors **(Fig. S1)**. Additionally, we show that the enhanced signal enables easy adjustment of the detection limit by controlling concentrations and reaction time, allowing for the detection of lead contamination at different regulatory limits in untreated spring water samples. Furthermore, we discuss to what extent aTFs cross-reactivity in an *in vitro* setting impacts sensor selectivity.

## 2. RESULTS and DISCUSSION

### 2.1. Design of a coupled nucleic acid lateral flow and *in vitro* transcription-based sensor

Successful cell-free biosensors have been developed based on IVT, utilizing fluorescence as the transduction method by leveraging the properties of RNA light up aptamers (Jung et al., 2022, 2020; Patino Diaz et al., 2022). However, there are still limitations in the achieved limit of detection (LoD) for certain species, such as lead (Ekas et al., 2024a; Jung et al., 2020; Li et al., 2025). To boost sensitivity while maintaining simplicity and compatibility with PoC we turned to a NALF assay, where detection is based on nucleic acid hybridization. We designed DNA oligonucleotides to directly detect IVT products using lateral flow assays. Our design involves two oligonucleotide probes (capture and reporter probes) which must simultaneously hybridize to a complementary sequence with fast kinetics (**Fig. S1**). The complementary sequence is encoded in the DNA template to be transcribed (**Fig. 1A**). We started by selecting the RNA output sequence. We decided to employ the IVT-compatible light-up aptamer 3WJdB (which contains two Broccoli units) and a transcript encoding a single Broccoli aptamer. This enables monitoring both the transcription kinetic parameters and the quantity of IVT products through fluorescence assays. It also facilitates direct comparison between the NALF sensor performance with fluorescence sensors reported in the literature. Despite its slow folding and high ratio of transcript per fluorescent molecule, the transcription efficiency of this aptamer is comparable to shorter RNA molecules frequently used in the literature with the additional advantage in the case of 3WJdB having multiple binding sites that can be recognized by NALF. Our hypothesis was that incorporating multiple binding sites within the RNA transcript could enhance the LFA signal through avidity effects.

**Fig. 1.**
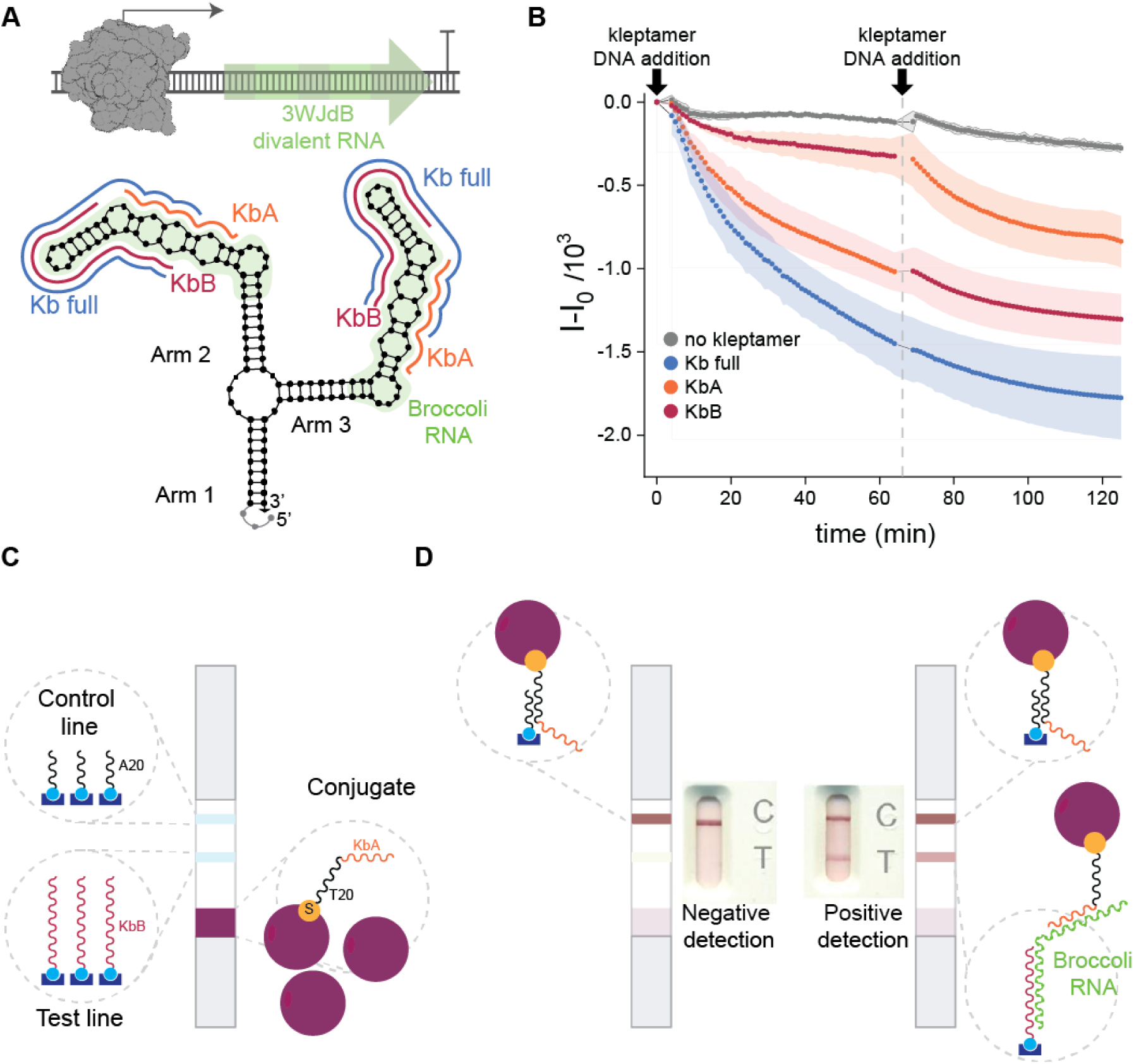
Design of a nucleic acid-based lateral flow (NALF) assay to detect IVT product. (A) Schematic representation of the IVT product, consisting of a divalent 3WJdB RNA and the designed kleptamers, which are complementary to the 3WJdB. (B) Kinetic experiments showing the decrease in fluorescence of the 3WJdB−DFHBI-1T complex formed after the IVT reaction, upon the addition of different kleptamer strands. The decrease in fluorescence is due to the hybridization of the kleptamers with the 3WJdB, which displaces the Broccoli aptamer target, the DFHBI-1T dye. (C) Schematic illustration of the designed lateral flow test. The lateral flow device contains a sample pad, a conjugate pad with gold nanoparticles conjugated with KbA sequence, a test line containing KbB kleptamer, a control line formed by A20 sequences, and an absorbent pad. (D) Representation of positive (equivalent to 5×10^−4^ MEF) and negative assay results. In the presence of the IVT product (positive), the 3WJdB is captured by KbB on the test line, while AuNPs containing KbA sequences hybridize with the aptamer, producing a colored line on the test line (T). In both cases, AuNPs are captured in the control line (C) using the T20 tail of the reporter sequence. A representative example of the visible signal for both positive and negative results is shown in the center.

Then, we designed the capture and reporter probe. Although 3WJdB and Broccoli aptamer are structured RNA sequences, antisense RNA nucleic acid sequences of the Broccoli aptamer, termed kleptamers, have been shown to disrupt its structure and impede the binding of the dye (DFHBI-1T), thereby reducing the fluorescence measured in the sample (Lloyd et al., 2018). Taking advantage of these results, IVT-NALF DNA probes were designed from the sequence either partially or fully complementary to the Broccoli aptamer. Moreover, to target the 3WJdB structure which contains two Broccoli aptamers along with the arms, we designed two new, longer DNA-kleptamers aimed at disrupting this structure. One was designed to disrupt the stem loop of the Broccoli aptamer, and the other one to bind the bulge region (which serves as the binding pocket for DFHBI-1T); these were named kleptamer KbB and KbA, respectively (**Fig. 1A and Table S1**). Additionally, a fully complementary sequence to the aptamer was studied (kleptamer KbFull).

The success of NALF assay is largely dependent on the kinetic properties of the recognition elements (Jauset-Rubio et al., 2016). Thus, we first aimed to identify kleptamers with the most effective hybridization process. This was achieved by a fluorescent assay that allowed assessment of the disruption of the Broccoli aptamer structure. In short, we added the DNA kleptamer to a solution containing the 3WJdB−DFHBI-1T complex and measured the fluorescence of the solution over time. The decrease in fluorescence correlates with the strand displacement reaction which allows us to follow the kinetic of hybridization reactions. It is worth noting that the concentrations of this solution-based assay were selected to allow comparison across different conditions and provide insight into the binding interactions, however the timing and concentration dynamics of the chromatographic process on the strip are so the assay can be completed in 15 min (*vide infra*).

Initial results showed that while a decrease in fluorescence intensity was observed at the beginning, fluorescence began to increase after a few minutes (**Fig. S2**). We hypothesized that a competition between the ongoing IVT reaction and strand displacement was occurring. To address this, we stopped the IVT reaction after 30 minutes, before adding the kleptamers, by heating the reaction to 65°C for 10 minutes to denature the T7 polymerase. **Fig. 1B** presents the kinetic measurements obtained after introducing different kleptamers to the heated IVT product. The continuous decrease of the fluorescence intensity obtained after ending the IVT reaction confirmed the need to stop the IVT reaction to observe strand displacement and opening of the 3WJdB structure. Additionally, we examined the effect of adding kleptamers KbA and KbB in different arrangements, each at 50 pmol. Interestingly, the decrease in fluorescence intensity is steeper when KbA is added compared to KbB. Moreover, the total decrease in intensity is greater when KbA is introduced first rather than KbB, even though the total amount of kleptamers is the same in both cases. This suggests that kleptamer KbA may have a greater disrupting effect. Finally, adding 100 pmol of KbFull, results in a slightly greater decrease in fluorescence intensity than adding KbA followed by KbB. Overall, all three kleptamers are capable of inducing the opening of the 3WJdB, as evidenced by a decrease in fluorescence, with KbA demonstrating a more pronounced effect than KbB.

Based on these kinetic measurements, we chose KbA as the reporter sequence and KbB as the capture sequence. Since the reporter sequence will be on the conjugate, it will interact with the IVT product first, ensuring the opening of the 3WJdB when the product reaches the test line and binds to the capture sequence. The capture sequences were modified with biotin at the 3’ end to enable their incorporation into the nitrocellulose membrane (test line) through avidin-biotin interactions. The reporter sequence was conjugated to AuNPs by modifying these sequences with a thiol group. Additionally, to consider the test valid it is necessary to capture AuNP conjugates in the control line. To achieve this, a 20-poly adenine (polyA) sequence was added to the control line, while a 20-poly thymine (polyT) sequence was attached to the 5’ end of the reporter sequence (see **Table S1**). This configuration ensures that the AuNPs conjugates are captured in the control line via polyA-polyT interaction. **Fig. 1 C and D** schematized this configuration.

### 2.2. NALF detection of IVT products allows signal intensification

Once the kinetics results allowed us to identify the reporter and capture sequences, we turned to assembling and evaluating the performance of the IVT-based NALF assay. Beyond adapting our previously developed immuno lateral flow strips protocols to prepare the NALF lateral flow strips (**Table S2**) (Peri Ibáñez et al., 2024), we optimized reporter and capture sequence immobilization. First, we incorporated the capture sequence onto the test line through avidin/biotin interaction. Different concentrations of the avidin and capture sequence were tested. A 1.5:1 ratio between avidin and capture sequence adding 10 pmol in the test line yielded the best results (**Table S3**). Second, different conjugation methods for the reporter sequence to the AuNPs were evaluated, including freeze-thaw(Liu and Liu, 2017a) and salt-aging(Liu and Liu, 2017b), along with various blocking conditions. Through determining the position of the LSPR (**Fig. S3**), the simplicity of the conjugation method, the yield, and the stability of the conjugate (**Table S4**), we conclude that the freeze-thaw method followed by a BSA/PEG blocking step is the best strategy. A representative final strip with the cassette is shown in **Fig. S4**. We then optimized the quenching conditions to avoid interference of an ongoing IVT reaction with the strand displacement (**Fig. S5**); in all the assays 5 mM EDTA was included in the LFA sample buffer where the IVT product is collected before addition on the strip. In summary, **Fig. 1D** presents the results of an IVT reaction (without regulation) on the optimized LFA strips, displaying an intense band at the test line in the presence of the IVT product, and no band in the negative control.

Our preliminary results showed that even undetectable fluorescence leads to a strong signal in our NALF-IVT test (**Fig. 1D**). The positive signal is derived from 5×10^−4^ MEF, three orders of magnitude lower than what can be detected with a handheld illuminator (Jung et al., 2020). Thus, we propose that our designed NALF with an AuNPs-colorimetric readout may have an enhanced sensitivity that can allow us to lower the LoD for IVT product detection which can further allow for a wider dynamic range for chemical contaminant detection (*vide infra*). To test our hypothesis of an avidity effect when 3WJdB is recognized by the sequence on the test line, increasing the LoD of the IVT-LFA, we compared the performance in the IVT-LFA of the two different templates for the IVT reaction, one containing the 3WJdB and the other containing just one Broccoli aptamer (**Fig. 2A** and **Table S1**). First, we ran the IVT reaction for each template and stopped the reaction at different time points. A 10% dPAGE analysis of the IVT products at different time points was performed to evaluate the amount of RNA generated in each case (**Fig. S6**). The intensity of the gel bands normalized by the number of nucleotides for both templates indicates that the amount of RNA produced is similar in both cases (**Fig. S6**). Additionally, the fluorescence of the IVT products complexed with DFHBI-1T is measured at the endpoint of the IVT reaction, followed by our NALF assay. In this regard, the Broccoli aptamer is not optimized for its use *in vitro* conditions due to poor folding efficiency, resulting in low fluorescent signal compared with a similar amount of 3WJdB (Alam et al., 2017; Autour et al., 2016). While a NALF assay does not require a fluorescent or folded probe to be used, it serves in assay optimization by allowing us to correlate the fluorescence intensity (standardized to known amounts of fluorescein (FITC), **Fig. S7**)) of the IVT product with the intensity of the test line (**Fig. 2C**). It is worth noting that 1μl of the IVT reaction is added to 99μl of sample buffer before running the NALF assay, corresponding to a 1:100 dilution. Also, we measured the fluorescence of the IVT product prior to the 1:100 dilution used for the LFA, making it easily detectable with a plate reader. Then, we consider this dilution to correlate the MEFs and NALF test line intensity. The first key observation is that the IVT reaction using the 3WJdB template generates a visible test line signal after just 5 minutes, whereas the Broccoli template requires 60 minutes to produce a visible signal (**Fig. 2B**), even though the amount of RNA produced is the same or slightly higher than the 3WJdB template (**Fig. S6**). These results confirm that the avidity effect enhances the limit of detection (LoD) in our IVT-NALF design. Furthermore, **Fig. 2C** demonstrates that with the 3WJdB template, a visible signal (test line intensity greater than 1000) is achieved at an equivalent 0.1 × 10⁻³ MEF. This indicates that the NALF assay has a detection capacity of three orders of magnitude higher than portable fluorescence detection (where 0.1 MEF leads to barely visible fluorescence) and ten-fold higher than a laboratory fluorometer (LOD = 1 × 10⁻³ MEF), while maintaining a PoC assay.

**Fig. 2.**
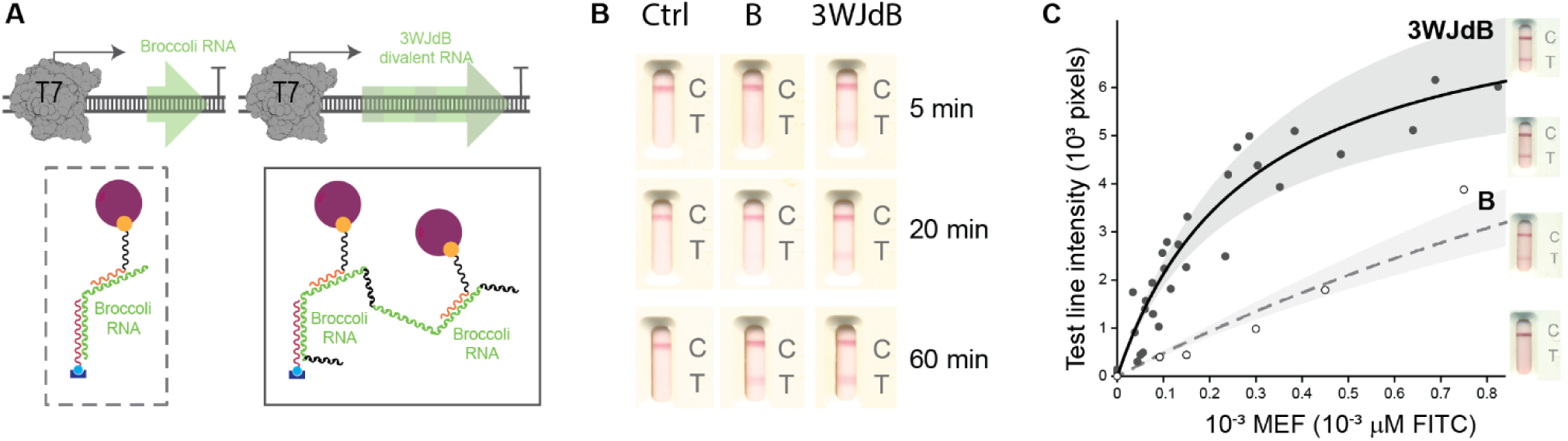
IVT products detected by a NALF assay show an amplification effect due to avidity. (A) Scheme comparing two tested IVT products: the Broccoli aptamer (left) and the three-way junction divalent Broccoli RNA, 3WJdB (right. The divalent Broccoli RNA can amplify the signal through avidity effects, as schematized. (B) NALF assay photos obtained after three different IVT reaction times (5, 20 and 60min) for a control condition (Ctrl, buffer only), Broccoli (B) and 3WJdB template. All the LFA assays scans are registered 20 min after the reaction is added to the strip. (C) Correlation between test line intensity of the NALF assays with the fluorescence intensity (MEF) from an IVT reaction. Black and white dots represent the results for 3WJdB and Broccoli IVT products, respectively. Each dot represents a single LFA with the sample obtained from a different IVT reaction, with 5 mM EDTA added in order to stop the progression of transcription. The test line intensity was measured as the number of pixels in the test line using photographs scanned with identical brightness and resolution. The LFA scans on the right show representative examples of the naked-eye signal, ranging from none, faint, intense, and very intense.

### 2.3. Improving the sensitivity of lead sensors by coupling to NALF-ROSALIND

Although cell-free IVT sensors show promise, detecting certain relevant metal ions, like Pb(II), at the low concentration required to meet regulatory standards still poses a considerable challenge. Therefore, we selected the CadC transcription factor-based ROSALIND system to evaluate our IVT-NALF assay to detect Pb(II). In the absence of Pb(II), the transcription factor CadC works as a repressor, preventing the IVT reaction from proceeding. In the presence of Pb(II), the binding of Pb(II) to CadC induces the release of CadC from the DNA, therefore the IVT reaction can proceed, and a signal is observed due to the formation of a 3WJdB(Jung et al., 2020). The signal enhancement in our IVT-NALF assay compared to portable fluorescence allows us to significantly decrease DNA and protein in the IVT reactions which can lead to a lower Pb(II) LoD (Li et al., 2025). To better understand how these concentration variations influence sensor performance, we used a single binding site model (one dimer to DNA) coupled with a monomer– dimer and metal-binding equilibrium (**Fig. 3A**) to fit the Pb(II) response curves at different CadC and DNA template concentrations (**Fig. 3B-D**). In this model, we utilized previously reported equilibrium constants for metal and DNA binding, when available, and fitted the response and obtained the remaining apparent constants from a global fit to all the protein, DNA, and metal concentration used (**Table S5**). Our data at 25 nM DNA template with 1.5 μM CadC (**Fig. 3B**, *black* dots) show that under these conditions the fluorescence significantly increases at Pb(II) concentration above 1 μM. Simulated curves corresponding to these DNA template concentrations at different CadC concentrations are also shown in **Fig. 3B** (*grey* lines). The simulations illustrate that decreasing the protein concentration may lead to a decrease in the LoD. We then performed the ROSALIND reducing the concentrations of the CadC repressor by 6-fold compared to previously reported work (Jung et al., 2020). Our results show that it is possible to decrease LoD by fivefold and twentyfold, albeit at the expense of a higher background signal (**Fig. 3C-D**). It should be noted, however, that the model fails to capture an increase in the unregulated signal observed when decreasing the protein concentration at constant DNA, suggesting that reduced CadC concentration can be beneficial for the signal at high Pb(II) (**Fig. S8**). These results show it is possible to tune the LoD by changing the ROSALIND condition. However, the possibility of translating this observation to a portable sensor is not direct because of the high background signal observed in the conditions where the LoD is lower, and the need of a highly sensitive detection method to detect low concentration of RNA output. Thus, leveraging the signal enhancement demonstrated by the IVT-NALF assay, we evaluated whether the NALF response could be tuned to exploit the lower LoDs observed at reduced protein concentration. We performed the ROSALIND-NALF assay using the ROSALIND conditions evaluated by fluorescence (2.5 nM and 25 nM DNA). To compare the response between different portable detection methods, the ROSALIND reactions were performed in the presence of the DFHBI-1T, thus the product was simultaneously detected using a portable fluorometer and with our NALF assay. With the naked eye, a handheld illuminator can detect 1 μM of Pb(II) using 25 nM of DNA template and 1.5 μM CadC, consistent with previously reported results (**Fig. 3 E**) (Jung et al., 2020). Under this condition, our NALF assay gives visible lines under all the tested concentrations, albeit a faint signal is also present at 0 μM Pb(II) (**Fig. 3E, S9A**). Thus, in this case NALF detection does not allow for sensor improvement. In contrast, our NALF assay exhibited ultrahigh sensitivity yielding markedly different results with other two ROSALIND conditions that impact the LoD (**Fig. 3 F-G, S9 B-C**).

**Fig. 3.**
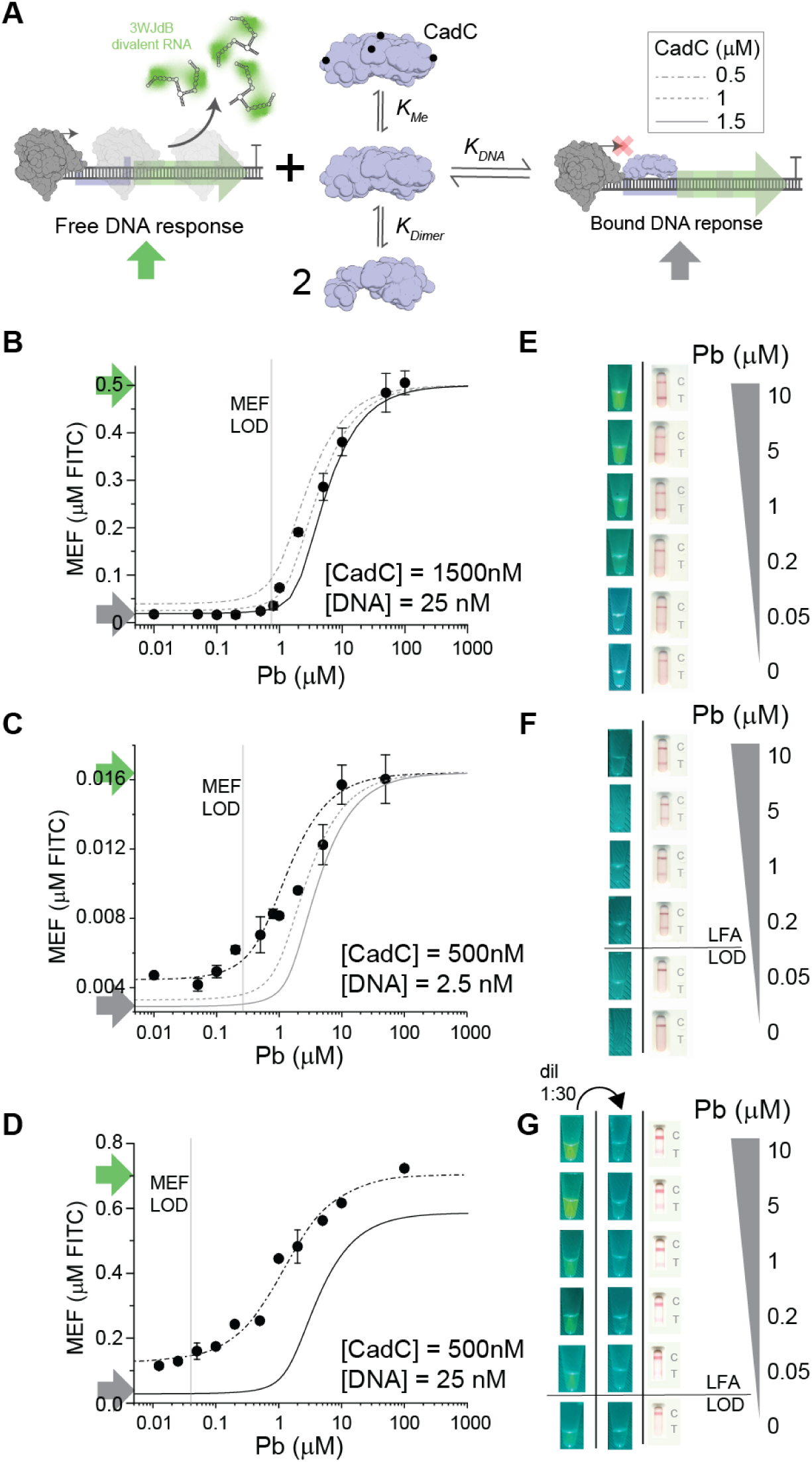
Pb(II) detection using ROSALIND-NALF. (A) Schematic representation of the equilibria involved in the IVT reaction regulated by the CadC repressor transcription factor. A model based on known equilibrium constants was developed to simulate Pb(II) response curves at varying concentrations of CadC and DNA template. (B) Standardized fluorescence (MEF) for different concentrations of Pb(II) in IVT assays (black dots) performed with 25 nM DNA template and 1.5 µM CadC, (C) with 2.5 nM DNA template and 0.5 µM CadC and (D) with 25 nM DNA template and 0.5 µM CadC. Lines represent simulated curves for various CadC concentrations at each DNA template concentration using the model described in (E) and (F) Images show the fluorescent and colorimetric signal from the portable fluorescence assay and the NALF assay at different Pb(II) concentrations. Reactions were stopped with the addition of 5mM EDTA at 20 minutes then a mixture of 1 μL of ROSALIND reaction with 99μL of NALF running buffer were added to test strips and test line intensities were measured. Reducing the DNA template concentration allows for a lower CadC concentration, enabling a lower limit of detection (LoD), though with increased background signal. By optimizing the conditions, an assay that generates a colorimetric signal only at Pb(II) concentrations above 0.2 µM can be achieved. (G) Images show the fluorescent and colorimetric signal from the portable fluorescence assay and the NALF assay at different Pb(II) concentrations. In this case, reactions were performed, stopped as previously described and then diluted in a 1:30 ratio prior to the 1:100 dilution in order to avoid the background signal observed in E. The reduction of CadC concentration while keeping a high DNA concentration resulted in a high fluorescent background, but with the dilution of the reactions we were able to obtain a lower LoD without a background signal in the NALF assay with colorimetric signal above 0.05µM Pb(II).

When the DNA template and CadC concentration were reduced to 2.5 nM and 0.5 μM, the NALF assay clearly detected 0.2 μM Pb(II), with no background signal (**Fig. 3F**, **S9B, S10**). These results contrast the lack of detectable fluorescence obtained in a handheld device (**Fig. 3F**), while agreeing with the fluorescence data acquired in the laboratory plate reader (**Fig. 3C**). Thus, NALF detection allows for the signal enhancement necessary for a portable test that meets the limit established by the Argentinian Food Code for drinking water (50 ppb, 0.2 μM) (“Codigo Alimentario Argentino (Argentine Food Code),” n.d.). On the other hand, reducing the CadC concentration to 0.5 μM while maintaining 25 nM DNA resulted in visible signals in the handheld illuminator at all tested Pb(II) concentrations, likely due to elevated background (**Fig. 3G**). Thus, taking advantage of the NALF signal enhancement, we performed the detection on diluted ROSALIND product, successfully detecting Pb(II) at concentrations corresponding to internationally accepted limits, such including the WHO guideline (10 ppb, 0.048 μM) (WHO, 2022) (**Fig 3G, S9C, S10**). Notably, under the same dilution conditions, no signal was detected using the handheld illuminator at any Pb(II) concentration.

Overall, these findings highlight that the enhanced sensitivity of the ROSALIND-NALF assay enables fine-tuning the LoD for Pb(II) by adjusting both the DNA template and CadC concentrations, meeting both local and global regulatory standards. This tunability is possible because the IVT-regulated reactions can be coupled with a NALF detection strategy that minimizes false positives from background signal while remaining highly sensitive to low levels of RNA output (**Fig. S10**).

Next, we investigated the potential of the ROSALIND-NALF assay for field-deployed applications, specifically addressing a critical water-quality issue in Argentina: lead contamination in the Matanza-Riachuelo basin. The governmental agency ACUMAR is in charge of the monitoring and sanitation of the basin. To this end, periodical spring water sampling campaigns are carried out in 84 spots spread in the basin (**Fig. 4A**). The samples are analyzed through a battery of standardized methods (Ambiental, 2022). In particular, total metal ions (Ni, Zn, Cr, Cd, Pb) are quantified with AAS (Standard Methods, n.d.). Moreover, ensuring the shelf-stability of the assay is critical for PoC applications. In this case, this involves lyophilizing the ROSALIND reactions for subsequent rehydration with the sample. To achieve this, we adapted the previously reported freeze-drying protocol to our laboratory setup and evaluated how lyophilization and storage impacted sensor performance (**Fig. S11A**). The established freeze-drying protocol and preservation conditions kept the reactions in stable working conditions for up to 42 days. Preserved reactions were rehydrated with untreated and lead spiked spring water samples from the basin to prove their compatibility with complex samples (**Fig. S11B**). Given that the regulatory limit for Pb(II) in Argentina is set at 0.2 μM, we selected conditions that produce an on–off signal at this threshold, leveraging the tunable limit of detection of our IVT-NALF assay. Here, we focused our attention on 7 samples taken at different points of the basin. Five of them did not contain detectable amounts of metal ions (supplementary information, **Table S6**). These samples were measured with the ROSALIND-NALF tests by rehydrating lyophilized ROSALIND reactions with the samples, showing negative results. Then, we spiked different amounts of Pb(II) to study the response of the sensor in the environmental spring water samples (**Fig. 4B**). We observed a strong correlation between the results in the buffer and with spring water samples (**Fig. 3F**). Furthermore, when Sample 7 was analyzed using our ROSALIND-NALF assay, a positive signal was observed on the test line, indicating that the sample contained Pb(II) at a concentration above 0.2 μM. This result is consistent with the value obtained using the standard AAS method, which reported 2.1 μM of Pb(II). These findings demonstrate that our assay can successfully detect Pb(II) in spring water samples. When Sample 6 was tested, a weaker positive signal appeared on the test line. According to AAS analysis, this sample contained 2.6 μM of Zn(II), suggesting that Zn(II) acts as an interfering ion detectable by our sensor, as expected based on the known selectivity profile of CadC.

**Fig. 4.**
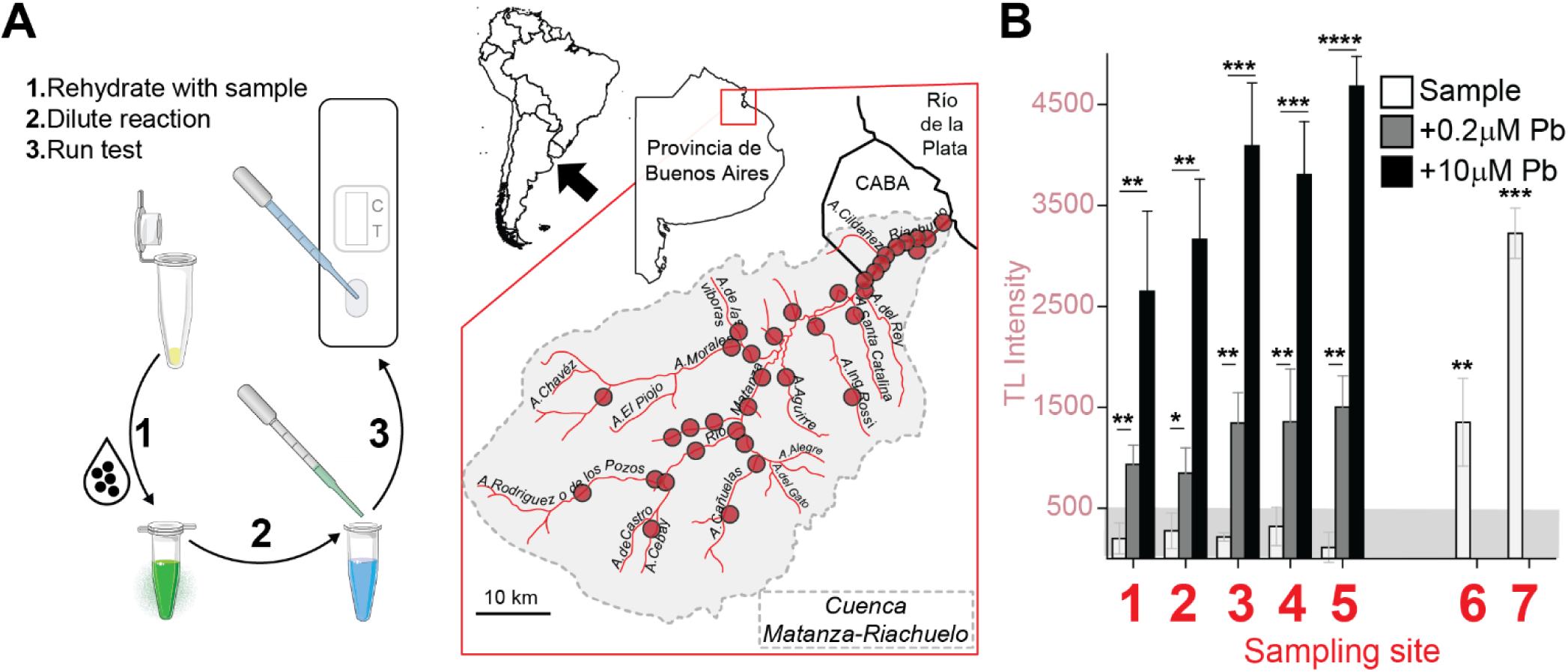
Pb(II) detection on spring water samples using ROSALIND-NALF. (A) ROSALIND reactions were rehydrated using samples collected from the Matanza Riachuelo basin at various sites from 84 samples obtained from anonymized sampling points represented on the map. (B) From these samples eight were selected for further testing (Table S6). The samples that did not contain Pb(II) were spiked with different Pb(II) concentrations, and measured before and after spiked with the NALF assay. n = 3 technical replicates (mean ± SD). The signal increment observed in spiked control samples were determined using a two-tailed Student’s t-test against the non-spiked condition, and their P value ranges are indicated with asterisks (* P=0.01-0.05, ** P=0.01-0.001, *** P=0.001-0.0001, *** P<0.0001). Samples’ 6 and 7 signals were compared individually against each control sample.

### 2.4. Evaluation of metalloregulators *in vitro* selectivity and field-testing applications

Cells buffer transition metal activity to ensure metalloproteins acquire their cognate metals under various conditions. This process depends on a complex interplay of factors, including the stability of metal-ligand complexes and the dynamics of the buffered metal pool, which is considered bioavailable and readily exchangeable (Capdevila et al., 2024, 2017; Clough et al., 2025; Foster et al., 2022; Lenner et al., 2025). Metalloregulators, which control the expression of metal transporters, are thought to be “tuned” to respond to a specific range of metal activity, preserving cellular metallostasis. However, this tuning is influenced by the intracellular milieu, creating a feedback loop in which the environment shapes and is shaped by metal selectivity. This interplay is further complicated by the varied inducer promiscuity of metallosensors, leading to crosstalk between regulons. Ultimately, the specificity of metalloregulation emerges from a delicate balance between sensor affinity, intracellular conditions, and the dynamics of the buffered metal concentrations. Moreover, many whole cell biosensors take advantage of selective accumulation of metals by expressing or deleting different transporter systems (Capdevila et al., 2017; Chen et al., 2023). Conversely, in cell-free conditions, where the chemical media is defined and metal ions are “available”, the selectivity of a metallosensor can only be defined by the intrinsic properties of its metal binding sites. In these conditions, non-cognate metals can drive the same *in vitro* allosteric response as the cognate metal thus affecting the selectivity of the sensor (Jordan et al., 2020). To study the degree to which crossreactivity affects ROSALIND reaction selectivity, we measured the response to different divalent metal cations in two closely related metalloregulators from the ArsR family (Diez et al., 2023), CadC and SmtB (**Fig. 5A, Fig. S12**). The signal after 20 minutes of reactions with 10μM of the corresponding ion shows that CadC responds to a number of divalent metals with a preference for Pb(II), that is improved when lowering the protein concentration (**Fig. 5A**, *top*). In contrast, under similar conditions SmtB responds only with Zn(II) (**Fig. 5A**, *bottom*). It is interesting to note that our SmtB data shows that SmtB can tightly repress the DNA template containing the CadC promoter sequence, *cadO* (**Table S5**). As CadC and SmtB belong to the same subgroup within the ArsR family (Diez et al., 2023), it is not unexpected that these transcriptional regulators would recognize the same DNA sequence. However, it is interesting to note that this observation may facilitate evaluating different metal specificities in related proteins by using a single DNA template as long as the aTF belongs to the same family or subfamily of proteins that recognize the same DNA sequence. This also has implications in our ability to build a NIMPLY logic gate between these closely related proteins (Jung et al., 2020) to prevent Zn(II) cross reactivity for CadC by harnessing SmtB specificity, as the transcription of a 3WjdB kleptamer is regulated by SmtB. However, it may not be possible using related aTF because of the promoter crosstalk. Altogether, our data suggest that both CadC and SmtB have different metal specificities with a preference for Pb(II) and Zn(II), respectively, over other transition metal ions even in the absence of the intracellular milieu.

**Fig. 5.**
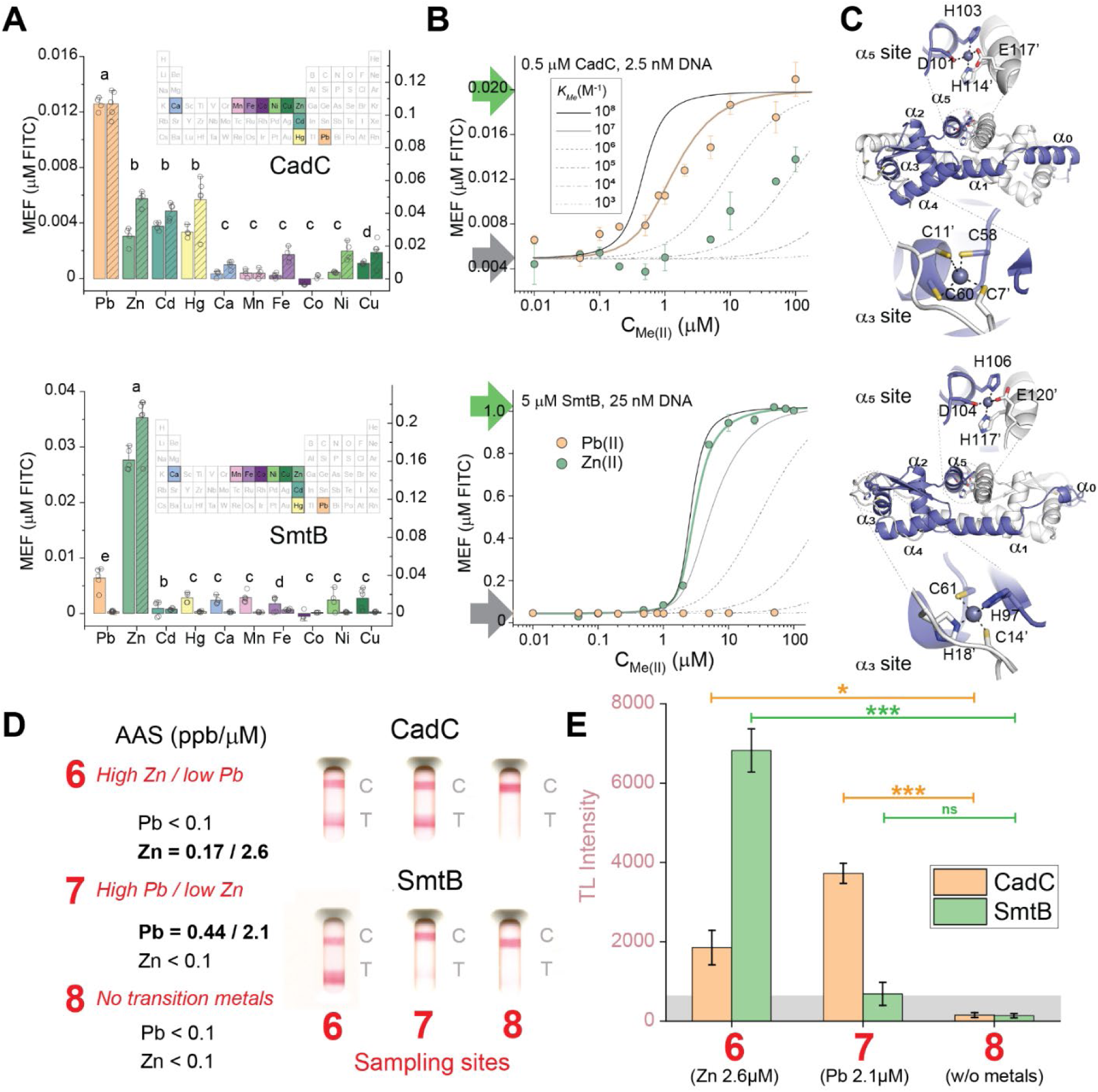
Selectivity of sensors based on metalloregulators and identification of high lead and zinc water samples. (A) Crossreactivity of CadC and SmtB-based ROSALIND sensors with different divalent metal ions. Standardized fluorescence (MEF) for 10 µM concentrations of divalent metal ions in IVT assays performed with 25 nM (dashed bars, right axis) and 2.5 nM DNA (full bars, left axis) template with *cadO* operator with 1.5 µM or 0.5 µM CadC (*top*), and 5 µM or 2 µM SmtB (*bottom*). Endpoint fluorescence was measured after 10 minutes. Groups sharing a letter are not significantly different according to Welch ANOVA with Games–Howell post-hoc test (p < 0.05), tests were performed for the conditions showed in the B panel. (B) Standardized fluorescence (MEF) for different concentrations of Pb(II)(*orange* dots) and Zn(II) (*green* dots) in IVT assays performed with 2.5 nM DNA template with *cadO* operator with 0.5 µM CadC (*top*), and 25 nM DNA template with *cadO* with 5 µM SmtB (*bottom*), endpoint fluorescence was measured at 10 and 60 minutes respectively. Lines represent simulated curves for various metal binding constants for CadC and SmtB. concentrations at each DNA template concentration. (C) Modeling of the zinc metalated forms of CadC (*top*) and SmtB (*bottom*) homodimer structures with each of the two protomers (*white* and *blue*). Close-ups of the two proposed metal binding sites with the ligands that are responsible for the different metal selectivity. These models have been deposited in Model Archive (https://modelarchive.org, with associated statistics). (D) Summary of the collected field samples with FAAS for metal concentration validation and the NALF assay results on those samples both with CadC (*top*) and SmtB (*bottom*) based sensors. (E) Intensity of NALF assays of the collected field samples. The error bars represent the standard deviation of assays performed in triplicate with the same field sample. The gray area marks the TL intensity threshold above which a naked-eye signal is visible. Signal levels measured for control water (8) and contaminated water samples (6 and 7) were compared using an unpaired two-tailed *t*-test, and their P value ranges are indicated with asterisks (ns P>0.01, * P=0.01-0.001, ** P=0.001-0.0001, *** P<0.0001).

To further characterize the extent of selectivity we analyzed the response curves of CadC and SmtB to varying concentrations of Pb(II) and Zn(II) under optimized conditions for ROSALIND-NALF assays (**Fig. 5B**). We focused our attention towards Zn interference as it is frequently detected in environmental samples at high and not toxic concentration (WHO, n.d.). These experimental results were fitted and compared with simulated curves generated using a previously described model fitted with different apparent metal-binding affinities (*K_met_*, **Table S5**). The selectivity of these sensors correlates with the difference in apparent *K_met_* values for the metals that are overall in agreement with the values reported in the literature. For SmtB, the *K_met_*for Zn(II) is more than three orders of magnitude larger than the corresponding to Pb(II) leading to vastly distinct response curves and LoDs. In contrast, CadC exhibits a *K_met_* for Pb(II) only two orders of magnitude larger than that for Zn(II), leading to more similar responses to both metals. These results suggest that for *in vitro* assays, *K_met_* is an appropriate indicator for metal selectivity in an *in vitro* setting.

To put our selectivity results in the context of what is known and what can be predicted based on the protein sequence, we constructed an Alphafold3 model of the Zn(II) bound form of CadC and SmtB homodimers (**Fig. 5C**). This model successfully recapitulated the two metal-binding sites reported for these proteins and conserved along the metal binding proteins in the ArsR family (Diez et al., 2023). The Zn(II) in α5 site is coordinated in both proteins by the same residues from both promoters: namely D101, H103, H114’ and E117’, in CadC, and D104, H106, H117’ and E120’, in SmtB. In contrast, the coordination of Zn(II) in α3 site differs significantly between CadC and SmtB. While CadC encodes for a distorted *S_4_*site which encompasses two Cys from each protomer, C58 and C60 in α3 and the N-termini C7í and C11í, SmtB harness a His_2_Cys_2_ site which likely a lower affinity for softer cations like Pb(II). These results are consistent with previously reported available crystal structures (Eicken et al., 2003; Ye et al., 2005), and with metal binding information of single point mutants (Busenlehner et al., 2002; Turner et al., 1996). The sole exception seems to be the α5 site in SmtB wild-type that may also involve C121 as shown by the presence of S coordination inferred from extended X-ray absorption fine structure (EXAFS) experiments (VanZile et al., 2000). It should be noted that these are Zn(II) bound structures and Pb(II) does not necessarily share the same coordination chemistry. Indeed, Cys reactivity experiments indicated that C11 is not involved in Pb(II) coordination (Apuy et al., 2004). Based on this data this site has been proposed to have pyramidal coordination as shown in PbrR (Huang et al., 2016). While the Zn(II) bound model cannot recapitulate the proposed Pb(II) binding site, it does predict a distorted *S_4_* where the pyramid is mainly defined by C11í, C58, and C60 while C7’ thiol is at a larger distance from the metal ion. Overall, the Alphafold3 model jointly with prior structural and functional information points to α3 being the main driver of the difference in selectivity. This is further supported by the observation of a lack of allosteric connection between the vestigial α5 site and DNA binding in *Sa*CadC (Busenlehner et al., 2002). Next, we examined if the selectivity difference between SmtB and CadC-based ROSALIND-NALF could be sufficient to distinguish high Zn(II) from high Pb(II) content in field samples. To address this, we took advantage of the spring water samples that have been fully characterized with different standardized protocols (See Materials and Methods, **Table S6**). First, we tested both SmtB and CadC based ROSALIND-NALF sensors with laboratory samples containing 10 µM Pb(II), Zn(II) and Zn(II)+Pb(II) to evaluate if the lateral flow readout introduced any interference in both assays. The results once again show good correlation between fluorescence (**Fig 5A**) and the intensity of the test line (**Fig. S**13). Then we turn to field samples 6 and 7 where Pb or Zn have been detected using AAS, as well as another sample where no transition metals were detected as a control. Neither sensor did not gave rise to false positives in the sample that did not contain either Pb or Zn despite the moderately high concentration of salts and other divalent ions in field samples (**Fig. 4B, 5D**, **S14, Table S6**). In these samples, only CadC sensors resulted in positive tests when the sample contained Pb. When Zn was found in the sample at a high concentration, SmtB sensors gave a strong signal and CadC sensors gave a positive signal as well (**Fig. 5D-E, S14**). While, as earlier discussed, CadC and SmtB sensors cannot be combined in a NIMPLY circuit since they recognize the same DNA operator, running additional tests with SmtB can prevent misinterpretation of CadC sensorsOverall, we can conclude that it is possible to identify Pb(II) contaminated spring water with CadC-based ROSALIND-NALF, and while the selectivity of CadC is not enough to prevent a false positive in field samples with Zn(II) concentration higher than 1 µM, running a test in parallel with SmtB may be sufficient to identify these samples (**Fig. 5D, S14**).

## 3. CONCLUSIONS

In this work, we reported the development of an IVT-NALF assay, demonstrating that by combining the amplification power of IVT with avidity-based nucleic acid detection, we can create rapid and ultra-sensitive point-of-care sensors capable of detecting relevant metal ion contaminants, such as Pb(II), at the low concentrations required. Our design takes advantage of avidity effects to reach a NALF assay with a detection capacity three orders of magnitude higher than portable fluorescence detection of a light-up aptamer **(Table S7)**. Furthermore, our results demonstrate that by adjusting concentrations and reaction times, the dynamic range can be easily modified to develop on-off sensors that detect contaminants at levels specified by various regulatory standards. For example, we developed Pb(II) sensors that activate at 0.2 µM (50 ppb), as required by Argentinian regulations, or at 0.048 µM (10 ppb), as required by the WHO. We then applied these sensors to detect Pb(II) in untreated spring water samples and successfully identified lead containing samples from the Argentinian Matanza-Riachuelo basin.

Moreover, we investigated how aTF–ligand crosstalk affects the selectivity of the ROSALIND-NALF assays. Typically, aTF selectivity is assessed under *in vivo* conditions to characterize their biological function. There aTF metal selectivity is attuned to the bioavailability of each metal ensuring proper protein metalation and tight regulation of cellular responses to either metal starvation or toxicity (Capdevila et al., 2024; Clough et al., 2025; Foster et al., 2022; Lenner et al., 2025; Osman et al., 2019). In this study, we evaluated the response of two closely related metalloregulators from the ArsR family, SmtB and CadC, to different divalent metal cations under *in vitro* conditions. It should be noted that while other speciation of the metal ions might be present in environmental water samples, the metalloregulators would respond primarily to oxidation status that would be more abundant in the cellular milieu. While there is a potential to harness the aTF selectivity to evaluate speciation, we focused our attention on ions that are more prevalent in spring water and are of bioinorganic interest, since they are part of the Irving-William series. We observed that SmtB exhibits specific selectivity for Zn(II), while CadC responds primarily to Pb(II) but also to Zn(II), Hg(II), and Cd(II). To minimize false positives, parallel reactions using both aTFs could be employed. Engineering the metal-binding site of CadC could further enhance selectivity, though achieving Pb in aTF selectivity remains challenging (Ekas et al., 2024a). Alternatively, introducing specific chelators that do not interfere with the transcriptional process or incorporating logic circuits such as NIMPLY(Jung et al., 2022, 2020) configurations could improve assay performance. For example, while SmtB and CadC share the same promoter, making them incompatible for logic circuits, aTFs with distinct operator architectures and desired metal selectivity, such as CsoR (Chang et al., 2014; Jung et al., 2020; Sakamoto et al., 2010), offer promising compatibility with circuit design. Furthermore, ROSALIND could be repurposed as a functional *in vitro* assay to directly assess aTF cross-reactivity and evaluate what are the ideal regulators for a specific inducer in a chemically defined media.

While our assay development focused on aTF-regulated IVT (ROSALIND) for detecting metal ion contaminants, other forms of IVT regulation could enable this highly sensitive IVT-NALF assay to detect a wide range of other analytes. For example, recent studies have employed nucleic acid structures, such as aptamers (Lee et al., 2024), CRISPR-Cas systems (Wang et al., 2025, 2022) or split T7 polymerases (McSweeney et al., 2025) to regulate IVT for the detection of proteins and nucleic acids. These regulatory mechanisms could expand the application of our assay beyond water contaminant monitoring to include diagnostic applications, even beyond exploiting the versatility provided by ROSALIND modularity that would allow exploring almost any relevant chemical contaminant (Capdevila et al., 2024; d’Oelsnitz et al., 2022; Ekas et al., 2024b).

## Acknowledgements

We thank members of the Capdevila and Peinetti laboratories for their valuable insights that contributed to this work, in particular we acknowledge Sofia Liuboschitz for her assistance in the initial efforts in establishing IVT reactions locally. We would also like to thank Prof. Julius B. Lucks and Dr. Yueyi Li from Northwestern University for their help, guidance and support in establishing cell-free biosensing in Argentina. We would also like to thank ACUMAR for the environmental samples and support (CONVE-2022-99798516), we acknowledge Matias Parra and Ignacio Satori for the agreement that made this study possible. M.V.D acknowledges the support of the UNU-BIOLAC research fellowship for enabling the visit to Prof. Fernán Federici at Pontificia Universidad Católica de Chile, which provided training in molecular biology techniques. Camila Bodden from Harvard University for her help with language revisions.

## AUTHOR CONTRIBUTIONS

M.V.D. (Conceptualization: Equal; Formal analysis: Equal; Investigation: Lead; Methodology: Lead; Writing – original draft: Equal; Writing review & editing: Lead). C.S. (Conceptualization: Equal; Formal analysis: Equal; Investigation: Lead; Methodology: Lead; Writing – original draft: Equal; Writing – review & editing: Supporting). F.A.R. (Investigation: Supporting; Visualization: Supporting; Writing review & editing: Supporting). G.G. (Investigation: Supporting; Methodology: Supporting; Resources: Equal). M.R. (Investigation: Supporting; Methodology: Supporting; Resources: Lead). D.A.C. (Conceptualization: Lead; Formal analysis: Supporting; Funding acquisition: Lead; Project administration: Lead; Supervision: Lead; Writing original draft: Lead; Writing – review & editing: Equal). A.S.P. (Conceptualization: Lead; Formal analysis: Supporting; Funding acquisition: Lead; Project administration: Lead; Supervision: Lead; Writing original draft: Lead; Writing – review & editing: Equal).

## SUPPORTING INFORMATION

Supporting information is available online.

## DATA AVAILABILITY

The data underlying this article will be shared on reasonable request to the corresponding author.

