## Supplementary material for "Lateral flow cell-free transcriptional assay for contaminant detection"

### Index

1. **Materials and Methods**
   - Plasmids and genetic parts assembly
   - Protein preparation
   - Plate reader quantification, fluorescein standardization and data fitting
   - IVT reactions and freeze drying
   - Freeze-drying
   - Municipal and surface water sampling and FAAS measurement
   - Gold nanoparticles and probe conjugation process
   - LFA Assay Preparation
   - LFT Evaluation and Sample Testing

2. **Supplementary Text**
   - **Supplementary Text 1.** Advancing Portable Biosensors for Affordable and Accessible Water Quality Monitoring
   - **Supplementary Text 2.** Cost analysis

3. **Supplementary Tables**
   - **Table S1.** Sequences used in the development and optimization of the lateral flow assay.

**Table S2.** Sets of parameters selected to build the strip.
**Table S3.** Optimization of the amount of capture sequence in the test line and the ratio avidin/biotinylated.
**Table S4.** Conjugation methods and parameters tested for the optimization of DNA-AuNP conjugate.
**Table S5.** Dimerization, DNA and metal binding affinities for CadC and SmtB
**Table S6.** Information of environmental superficial water samples.
**Table S7.** Comparison between Pb detection methods.

4. **Supplementary Figures**

**Figure S1.** Schematic representation of the ROSALIND-NALF assay.
**Figure S2.** Disruption of an IVT reaction with kleptamers, without stopping transcription.
**Figure S3.** UV-vis spectra of citrate-capped AuNPs and DNA-AuNP conjugates synthesized via different methods.
**Figure S4.** Representative strip showing the full assembly of all its components.
**Figure S5.** Effect of the addition of different concentrations of EDTA to the IVT reaction product.
**Figure S6.** dPAGE analysis of the IVT products at various reaction times for two different DNA templates.
**Figure S7.** FITC calibration.
**Figure S8.** MEF measurements for IVT reactions performed at 25nM DNA and different concentrations of CadC.
**Figure S9**. Replicates of the ROSALIND-NALF Pb(II) detection assays shown in Figure 3.
**Figure S10**. Detection limit determination at 0.5 µM CadC and different DNA concentration.
**Figure S11.** Stability of lyophilized ROSALIND reactions and monitoring results from water samples from the Matanza Riachuelo basin.
**Figure S12.** Transcription Factors purification quality controls.
**Figure S13.** Comparison of test line intensities for CadC- and SmtB-based ROSALIND-NALF sensors in the presence of high concentration of divalent cations.
**Figure S14.** Replicates of the ROSALIND-NALF Pb(II) detection assays results shown in Figure 5D

### Materials and Methods

#### Plasmids and genetic parts assembly

Overexpression plasmids for *Sa*CadC and *Sy*SmtB in pET3a backbones were obtained as previously described (Busenlehner et al., 2001; Kar et al., 1997). Genetic parts required for ROSALIND reactions were previously described (**Table S1**) (Jung et al., 2020). Modified Oligonucleotides sequences were purchased from Integrated DNA Technologies.

Transcription templates were generated by PCR amplification of the plasmid pJBL729 (https://www.addgene.org/140399/) with the primers underlined in the transcription template in Table S1. Amplified templates were then purified and verified for the presence of a single DNA band of expected size on a 2% Trisacetate–EDTA (TAE)–agarose gel. Concentrations were determined using a Nanodrop.

#### Protein preparation

*Sa*CadC and *Sy*SmtB were expressed and purified as previously described (Busenlehner et al., 2001). Plasmids were transformed into the Bl21(DE3)* *E. coli* strain. Cell cultures (1 L) were grown in Luria broth at 37 °C, induced with 1 mM of isopropyl-β-D-thiogalactoside at 0.6OD and induction occurred at 37 °C for 3h. Cultures were then pelleted by centrifugation and resuspended in degassed lysis buffer (25 mM 2-(N-morpholino)ethanesulfonic acid (MES) pH 6, 750 mM NaCl, 2 mM Tris(2-carboxyethyl)phosphine (TCEP), 1mM Ethylenediaminetetraacetic acid (EDTA), 0.1mM phenylmethylsulfonyl fluoride (PMSF)). Resuspended cells were then lysed on ice through ultrasonication, and insoluble materials were removed by centrifugation. The supernatant was treated with 0.015 %V/V of poly(ethyleneimine) solution. The suspension was centrifuged, and the supernatant was taken to 70% ammonium sulfate. The suspension was centrifuged and the pellet resuspended in a lysis buffer with 75mM NaCl for subsequent dialysis with the same buffer in a 10kDa dialysis bag overnight. The dialysis product was centrifuged, and the supernatant loaded into a SP cation exchange column to be eluted with a gradient of lysis buffer. The fractions containing protein were pooled and treated for 16 hours with Tobacco Etch Virus (TEV) protease, then loaded into a Superdex 75 size exclusion column equilibrated with S75 buffer (25mM Tris,

200mM NaCl, pH 8, degassed, 2mM TCEP). Fractions corresponding with the protein's weight were pooled and concentrated. Protein concentrations were measured via 280nm absorbance, and protein purity and size were verified via SDS-PAGE. Purified proteins were stored at -80 °C as single use aliquots.

T7 RNAP plasmid was first transformed into Bl21(DE3)* *E. coli* strain. Cell cultures (1 L) were grown in Luria broth at 37 °C, induced with 1 mM IPTG at 0.6OD and let induction occur at 37 °C for 3h. Cultures were then pelleted by centrifugation and resuspended in lysis buffer (20mM TRIS pH=8, 500mM NaCl, 5% v/v Tween 20, 5mM imidazole and 0.1mM PMSF). Lysis was done via ultrasonication and then centrifuged. The clarified supernatant was loaded into a histrap HP column and eluted with a gradient against 20mM TRIS pH=8, 500mM NaCl, 500mM Imidazole. The fractions containing protein were pooled, concentrated and buffer exchanged with a centricon with 2xStorage buffer (100mM TRIS pH=7.6, 300mM NaCl, 0.2mM EDTA, 2mM dithiothreitol (DTT)) aiming to obtain a final concentration of 100µM of protein. This stock waslater diluted with glycerol to 50% and stored in -80°C as single use aliquots ensuring that fresh reducing agent is either present of added before reaction. Protein concentrations were measured via 280nm absorbance, and protein purity and size were verified via SDS-PAGE.

**Plate reader quantification, fluorescein standardization and data fitting**

A National Institute of Standards and Technology (NIST) traceable standard (Invitrogen, catalog no. F36915) was used to convert arbitrary fluorescence measurements to molecules of equivalent fluorescence (MEF). Serial dilutions from a 5-µM stock were prepared in a 100 mM sodium borate buffer at pH 9.5, including a 100-mM sodium borate buffer blank (total of 12 samples). The samples were prepared in technical and experimental triplicates (12 samples × 3 replicates = 36 samples total), and fluorescence values were read at an excitation wavelength of 487 nm and emission wavelength of 510 nm for Three Way Junction dimeric Broccoli (3WJdB)-activated fluorescence on a plate reader (Varioskan LUX). Linear regression was then performed for concentrations within the linear range of fluorescence (0–0.625 µM fluorescein isothiocyanate (FITC)) between the measured fluorescence values in arbitrary units and the concentration of FITC for identifying the conversion factor.

The fluorescence data from different times after transcription started were fit to a single binding site model (one dimer to DNA) coupled to a monomer–dimer equilibrium and metal binding equilibrium using Dynafit and previously reported binding constants. The schematics of the model used are presented in Figure 3A and the binding constants obtained are summarized in **Table S5** (Kuzmič, 1996).

**IVT reactions**

Homemade IVT reactions were set up following conditions adapted from the established ROSALIND protocols (Jung et al., 2020; Jung et al., 2022; Kartje et al., 2021). To ensure reproducible results the quality of the DNA template was verified as indicated above. rNTP solutions were prepared from solid stocks adjusting the pH to 7.0, preventing acidification of the reaction mix or hydrolysis risks associated with excessively high pH. For optimal performance, single-use aliquots of ROSALIND components, DNA template, rNTPs, IVT buffer (40 mM Tris-HCl pH 8, 8 mM $MgCl_2$, 10 mM DTT, 20 mM NaCl, and 2 mM spermidine), and aTFs were prepared, to prevent signal degradation caused by repeated freeze–thaw cycles. Unregulated reactions were set up by adding the following components listed at their final concentration in order: IVT buffer, 0,02mM (5*Z*)-5-[(3,5-Difluoro-4-hydroxyphenyl)methylene]-3,5-dihydro-2,3-dimethyl-4*H*-imidazol-4-one (DFHBI-1T), 11.4 mM Tris-buffered nucleotide triphosphates (2.85mM each), pH 7.0, 0.015 U thermostable inorganic pyrophosphatase DNA transcription template and MilliQ $H_2O$ to a total volume of 18 µl. Regulated IVT reactions additionally included a purified aTF and a ligand at the indicated concentrations and were equilibrated at 37 °C for 15 min. Immediately before plate reader measurements, 2µL of 50µM T7 RNAP was added to the reaction. Reactions were then characterized on a plate reader as described under Plate reader quantification and MEF standardization.

**Freeze-drying**

Lyophilization was done adapting established protocols (Jung et al., 2020; Jung et al., 2022). Briefly, before lyophilization, PCR tube caps were punctured with a syringe needle to create three holes. Lyophilization of IVT reactions was then performed by assembling the components of regulated IVT adding 50 mM trehalose and 250 mM d-mannitol in a reaction mix. This reaction mix was distributed in 20µL aliquots in pre-chilled tubes on an aluminum rack on ice. The racks were then submerged in liquid nitrogen and then transferred to a bench freeze dryer for 3 hours of freeze-drying with a condenser temperature of −85 °C and 0.04 mbar of pressure. Freeze-dried reactions were packaged in vacuum-sealed bags with a desiccant and oxygen absorber. The packets were stored in dark containers until rehydration.

**Municipal and surface water sampling and FAAS measurement**

Samples were collected simultaneously in March 2022, from the 53 of 130 sampling stations, by filling a 4L container from the surface water body, thereafter, taken to the laboratory in plastic bottles. pH, conductivity and total Ca/Mg, Ni, Zn, Cr, Cd and Pb concentration were determined in each sample pool. Briefly, the samples were digested with the addition of concentrated nitric acid until a light colored, clear solution was obtained. Then, heavy metal concentrations were measured using air/acetylene flameless atomic absorption with an iCE 3300 (Thermo Scientific) (Standard Methods, n.d., n.d.), and Ca/Mg were measured by an EDTA titrimetric method (Standard Methods, n.d.). From these 53 samples studied by fluorescence presented in **Fig. S11B**, eight samples were selected for further analysis and, therefore, all their parameters are presented in **Table S6**. Additional information about these sampling sites can be found in ACUMAR website (*ACUMAR | Autoridad Cuenca Matanza Riachuelo*, n.d.).

**Gold nanoparticles and probe conjugation process**

Gold nanoparticles (AuNPs) with a diameter of 20 nm were synthesized using a modified Turkevich protocol. Briefly, glassware was prepared by repeatedly washing with aqua regia (HCl + $HNO_3$, 3:1 ratio) followed by thoroughly rinsing with ultrapure water. A solution of $HAuCl_4$ (0.01%) was brought to boil under constant stirring and at this point a freshly prepared 1% sodium citrate solution was rapidly added. The solution was maintained at boiling for 10 minutes, then transferred to a cooling bath until it reached room temperature. These 20 nm AuNPs served as seeds for further growth. To grow the nanoparticles to 30 nm, a diluted AuNP seed solution was mixed with 1% sodium citrate and brought to a boil under constant stirring. Once boiling, $HAuCl_4$ (0.2%) was added, with further additions at 5 and 7 minutes after the initial addition. The solution was kept at boiling throughout and then cooled in a water bath to room temperature. The final suspension was stored at 4°C in the dark until use. The synthesized AuNPs were characterized by UV-Vis spectroscopy and scanning electron microscopy (SEM) (Peri Ibáñez et al., 2024).

To prepare the DNA-AuNPs conjugates, thiolated DNA probes were incubated with 20 mM TCEP at room temperature for 1 hour. After treatment, two conjugation methods were followed and compared. For the freeze-thaw protocol, the activated DNA probes were added to the 30 nm AuNP suspension to a final concentration of 4 µM. Conjugation was carried out using a single freeze-thaw cycle, wherein the mixture was frozen for 2 hours at -20°C and then thawed (Liu and Liu, 2017a). DNA was added dropwise to the AuNP solution during this process. To block any remaining free sites on the AuNP surface and enhance stability, bovine serum albumin (BSA, 10%) and polyethylene glycol (PEG-20000, 1%) were added. For the salt aging protocol, the activated DNA probes were added in excess to the 30 nm AuNP suspension to a final concentration of 7 µM (Liu and Lu, 2006). The mixture was incubated overnight at room temperature, protected from light. After incubation, 1 M NaCl was added dropwise with gentle shaking to gradually increase the ionic strength, reaching a final NaCl concentration of 150 mM. Drops were added every 20 minutes. The sample was then incubated overnight at 4°C while kept away from light. Following the salt stabilization step, mercaptohexanol (MCH) was freshly prepared and added to a final concentration of 1 µM. The solution was incubated at room temperature for 1 hour to complete the functionalization process. Both conjugated AuNPs suspensions were centrifuged at 8,000 RPM for 15 minutes at 4°C, and the supernatant was removed. A 1:200 dilution was analyzed by UV-Vis spectroscopy to evaluate the localized surface plasmon resonance(LSPR). The conjugated AuNPs were then resuspended in a pH-adjusted buffer (pH 7.5) to achieve the desired optical density (OD).

**LFA Assay Preparation**

To prepare the DNA-labeled test and control lines, 10 pmol of each DNA biotinylated probe were incubated with avidin for 1 hour at room temperature. Complexes were prepared at a concentration of 10 pmol/µL in phosphate-buffered saline (PBS) and were designated for the test line (KbB) and control line (A20). Nitrocellulose membranes (6 × 30 cm) were mounted onto a backing card, and the probes were dispensed using BioDot BioJet Quanti dispensers. The test line was dispensed at a rate of 3 µL/cm, while the control line was dispensed at 2 µL/cm. The previously prepared AuNP-DNA conjugate suspension was applied to a glass fiber pad using the AirJet feature on the BioDot dispense system. The sample pad was treated with a solution of PBS with 0.5% Tween-20. After dispensing, the membrane, the conjugate pad and the sample pad were dried in an air-dry oven at 37°C for 1 hour. Once all pads were prepared, they were assembled onto the backing card with a manual lamination system, overlapping each component to create a continuous flow path. The assembled membranes were then cut into 3.8 mm strips using a guillotine cutter to fit standard plastic cassettes.

**LFT Evaluation and Sample Testing**

Prepared test strips were stored in a low-humidity environment until they were assembled into cassettes. For testing, a running buffer composed of PBS with 0.5% Tween-20 was prepared. After performing the IVT reaction as described previously, the reaction was halted either by heating at 75°C for 20 minutes or by adding 2 µL of 50 mM EDTA to the samples. Our previous results indicated the need to halt the IVT reaction before any strand displacement occurs. To avoid heating, as we intended for Point-of-Care (PoC) settings, we explored the use of EDTA as an alternative method. EDTA is known to bind metal ions that are necessary for the IVT reaction to proceed. We tested various concentrations of EDTA**.** The increase in fluorescence after the addition of EDTA indicates that at concentrations lower than 1 mM, the IVT reaction continues. In contrast, at concentrations higher than 10 mM, the sudden decrease in fluorescence suggests that EDTA might be interfering with Broccoli aptamer correct conformation. Consequently, we found that adding 2 to 5 mM EDTA is an effective method to stop the IVT reaction and is more suitable for PoC applications. Thus, the addition of EDTA was incorporated into the assay by adding it directly to the IVT reaction product to halt the reaction before applying the sample to the strip.

For the assay, 10 µL of the running buffer was mixed with 1 to 5 µL of the IVT reaction product, depending on the experimental conditions. The mixture was then diluted with IVT buffer to a total volume of 100 µL. For negative control samples, IVT buffer was used in place of the IVT reaction product. This prepared solution was applied dropwise onto the sample pad window of each cassette. After a 20-minute incubation, the strips were scanned with consistent resolution settings, and images were analyzed using ImageJ software. The test line intensity was measured by selecting a specific Region of Interest (ROI) and quantifying the number of pixels. The intensity of the control line served as an internal control for each assay.

### Supplementary Text

**Supplementary text 1.** Advancing Portable Biosensors for Affordable and Accessible Water Quality Monitoring

Ensuring access to reliable information on drinking water safety remains an unresolved challenge. In many cases, decision making about water relies exclusively on perceptions and past experiences.(Miller et al., 2024) This situation is accentuated in regions of low- and middle-income countries as water contamination due to pollution poses significant risks to public health and monitoring infrastructure is often inadequate.(WHO/UNICEF Joint Monitoring Programme for Water Supply, Sanitation and Hygiene (JMP), 2021) Given the high burden of disease associated with contaminated water consumption, simple and affordable contaminant detection techniques are required.(Lee et al., 2023; PureEarth, n.d.; WHO, n.d.) In this context, portable biosensors have emerged as a promising solution for rapid, on-site monitoring of chemical contaminants in water, ultimately enabling testing water quality in settings with limited infrastructure, resources, and regular monitoring capabilities.(Chen et al., 2023; Huang et al., 2023; Li et al., 2023; Thavarajah et al., 2020; Yang et al., 2021) However, the application of these devices in communal settings has largely been limited to initiatives from developed countries and often fails to achieve long-term adoption by local communities.(Nadra, 2024; Thavarajah et al., 2023) A key bottleneck has been the development of optimal and cost-effective portable devices that allow simple result interpretation. The local development and production of biosensor reagents and devices appears as a critical step to enable affordability and accessibility of these technologies across vulnerable regions.(Arce et al., 2021; Bhamla et al., 2017; Cerda et al., 2024; Peri Ibáñez et al., 2024) Here, we address these challenges in the context of a noteworthy environmental challenge in Latin America which is lead monitoring in the Matanza Riachuelo basin. Since 2006, in response to evidence of lead contamination, local authorities have been building monitoring capabilities around this and other chemical contaminants in a basin that spans 2240 $km^2$ and is inhabited by 4.7 million people.(Olejarczyk, 2023; Pasqualini, 2019; Tripodi et al., 2023) Biosensors can play a crucial role in advancing this monitoring effort by enabling simple and affordable identification of sites with lead concentration exceeding regulatory limits. We propose that combining better readout strategies with advanced biosensing technologies alongside local production and distribution networks, biosensing innovations could achieve the promise of improving water safety across vulnerable regions.

**Supplementary Text 2.** Cost Analysis

The cost of individual ROSALIND reactions is primarily driven by the DFHBI-1T dye, at approximately US$0.35 per reaction. Additional reagents, depending on production and sourcing, can bring the total to 0.67 USD(Jung et al., 2020). Coupling ROSALIND reactions with different readouts impacts both cost and performance. For example, an amplification scheme based on modified oligos achieves a 10-fold reduction in LOD, with an associated cost increase to US$4.26 per reaction(Li et al., 2025). Lateral Flow Assay manufacturing costs are particularly sensitive to production scale: large-scale antibody-based devices cost around US$0.22 per test where more than half comes from the antibody cost, increasing to as much as US$3 for smaller-scale production runs(Rosen, 2009; Yetisen et al., 2013). In this context, the use of short DNA sequences as recognition elements reduces overall cost due to their lower production expense compared with monoclonal antibodies(Pan et al., 2018).

**Table S1.** Sequences used in the development and optimization of the lateral flow assay.

| Name | Full Sequence, 5' to 3' |
|---|---|
| **3WJdB IVT template** | GCGGATAACAATTTCACACAGGAAACAGCTATGACCATGATTACGCCAAGCTTGCATGCCTGCAGGTCGACTCTAGATAATACGACTCACTATAGGAGG**CTCAAATAAATATTTGAATGAA**CCCACATACTCTGATGATCCGAGA**CGGTCGGGTCCAGATATTCGT**ATCTG**TCGAGTAGAGTGTGGGCTC**GGATCATTCATGGCAAGAGA**CGGTCGGGTCCAGATATTCGT**ATCTG**TCGAGTAGAGTGTGGGCTC**TTGCCATGTGTATGTGGGTAGCATAACCCCTTGGGGCCTCTAAACGGGTCTTGAGGGGTTTTTTG |
| **Broccoli aptamer IVT template** | GCGGATAACAATTTCACACAGGAAACAGCTATGACCATGATTACGCCAAGCTTGCATGCCTGCAGGTCGACTCTAGATAATACGACTCACTATAGGAGG**CTCAAATAAATATTTGAATGAA**CCCACATACTCTGATGATCCGAGA**CGGTCGGGTCCAGATATTCGT**ATCTG**TCGAGTAGAGTGTGGGCTC**TAGCATAACCCCTTGG |

| | |
|---|---|
| | GGCCTCTAAACGGGTCTTGAGGGGTTTTTTG |
| **Kleptamer_KbA** | GG**GAGCCCACACTCTACTCGA** |
| **Kleptamer_KbB** | **ACGAATATCTGGACCCGACCG** |
| **Kleptamer_KbFull** | GG**GAGCCCACACTCTACTCG**ACAGAT**ACGAATATCTGGACCCGACCG**TCTCCC |
| **Reporter sequence (conjugate to AuNPs)** | /5ThioMC6-D/TTTTTTTTTTTTTTTTTTTTGG**GAGCCCACACTCTACTCGA** |
| **Capture sequence (test line)** | **ACGAATATCTGGACCCGACCG**TCTCCC/3Bio/ |
| **Control line sequence** | /5Biosg/AAAAAAAAAAAAAAAAAAAA |

**Table S2.** Sets of parameters selected to build the strip.

| LFT element | Parameter | Selected variables |
|---|---|---|
| Sample | Target to be detected | *in-vitro* transcription product |
| | Sample buffer composition | PBS w/Tween-20 0.5% |
| | Sample pad pre-treatment | PBS w/Tween-20 0.5% |
| Membrane | Test line | Capture DNA sequence |
| | Test Line dispensing buffer | PBS pH 7.4 |
| | Control line | Control line DNA sequence |
| | Nitrocellulose membrane | Hi-Flow 135 (Merck) |
| | Blocking | None/PBS+BSA 2% |
| Conjugate | Sequence | Bait DNA sequence |
| | Conjugation pH | pH 7.5 |
| | Sequence concentration | 4 µM |
| | Blocking | BSA and PEG 20000 |
| | OD | 5 |
| Pads | Sample | CFSP238000 (Merck) |
| | Conjugate | GFDX203000 (Merck) |
| | Wicking | CFSP223000 (Merck) |

**Table S3.** Optimization of the amount of capture sequence in the test line and the ratio avidin/biotinylated sequence using a half-strip assay. The best condition is highlighted in bold.

| Sequence amount (control and test lines) | Avidin-HRP amount | Strip |
|---|---|---|
| 5 pmol | 10 pmol | 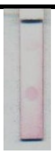 |
| 10 pmol | 5 pmol | 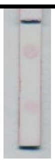 |
| 10 pmol | 10 pmol | 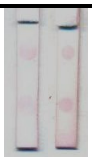 |
| **10 pmol** | **15 pmol** | 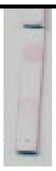 |
| 10 pmol | 20 pmol | 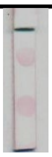 |
| 20 pmol | 20 pmol | 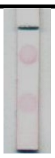 |

**Table S4.** Different conjugation methods and parameters tested for the optimization of DNA-AuNP conjugate.

| Conjugation method | Blocking agents | Volume obtained (µL) | Yield (%) |
|---|---|---|---|
| Salt-aging | MCH 1,12 µM | 25 | 48,3 |
| Freeze-thaw | BSA 3,5% + PEG 2000 0,2% | 150 | 79,2 |
| Freeze-thaw | None | 60 | 75,0 |

**Table S5.** Dimerization, DNA and metal binding affinities for CadC and SmtB

| | $K_1$ [x $10^6$ $M^{-1}$][a] | $K_2$ [x $10^6$ $M^{-1}$][b] | $K_3$ [x $10^6$ $M^{-1}$][b] |
|---|---|---|---|
| | ***dimerization*** | ***metal binding*** | ***DNA***[c] |
| **CadC** | 3 | Pb) 10 ± 4 ($n$ = 5)[d]<br>Zn) < 1 | 16 ± 7 ($n$ = 5)[f] |
| **SmtB** | 0.7 | Pb) < 0.01<br>Zn) 40 ± 20 ($n$ = 2)[e] | 12 ± 4 ($n$ = 2)[g] |

[a] The dimerization binding constants were fixed to the values reported in the literature for the apoproteins.(Busenlehner et al., 2001; Kar et al., 1997) [b] Experimental conditions: 40mM Tris pH=8, 8mM $MgCl_2$, 10mM DTT, 20mM NaCl, 2mM Spermidine. $K_a$ and errors were obtained from the global fit of all replicates using Dynafit software. [c] All experiments were performed with a template encoding for the CadC responsive operator (ctcaaataaatatttgaatgaa). The full sequence is reported in Table S1. [d] The lead-binding constant reported for this protein is 1.1 x $10^6$ $M^{-1}$. [e] The zinc-binding constant reported for this protein is $10^{10}$ $M^{-1}$. [f] The DNA-binding constant reported for this protein and the CadC responsive operator (t*ataatacactcaaataaatatttgaatgaagatg) is 1.1 x $10^9$ $M^{-1}$ in 5mM Mes, 400 mM NaCl, 1 mM DTT, 50 mM EDTA, pH 7.0 , 25°C.(Busenlehner et al., 2002) [g] The DNA-binding constant has not been

reported for this protein and any DNA sequence, however, EMSA experiments reported specific recognition of an SmtB responsive operator (aacacatgaacagttattcagatatt).(Turner, 1996)

**Table S6.** Information of environmental superficial water samples

| | | | Total | | | | | | | | | | |
|---|---|---|---|---|---|---|---|---|---|---|---|---|---|
| | **pH** | **Conductivity** | **Ca/Mg** | **Ni** | | **Zn** | | **Cr** | | **Cd** | | **Pb** | |
| | | µS/cm | ppm | ppm | µM | ppm | µM | ppm | µM | ppm | µM | ppm | µM |
| **1** | 7.24 | 397 | 92,6 | <0.03 | <0.5 | <0.01 | <0.15 | <0.06 | <1.15 | <0.006 | <0.05 | <0.03 | <0.14 |
| **2** | 7.99 | 1614 | 188,7 | <0.03 | <0.5 | <0.01 | <0.15 | <0.06 | <1.15 | <0.006 | <0.05 | <0.03 | <0.14 |
| **3** | 7.99 | 1614 | 200,9 | <0.03 | <0.5 | <0.01 | <0.15 | <0.06 | <1.15 | <0.006 | <0.05 | <0.03 | <0.14 |
| **4** | 7.66 | 956 | 168,4 | <0.03 | <0.5 | <0.01 | <0.15 | <0.06 | <1.15 | <0.006 | <0.05 | <0.03 | <0.14 |
| **5** | 7.96 | 890 | 185,5 | <0.03 | <0.5 | <0.01 | <0.15 | <0.06 | <1.15 | <0.006 | <0.05 | <0.03 | <0.14 |
| **6** | 8.11 | 1716 | 187,3 | <0.03 | <0.5 | <0.01 | <0.15 | <0.06 | <1.15 | <0.006 | <0.05 | **0.44** | **2.1** |
| **7** | 7.26 | 1760 | 228,4 | <0.03 | <0.5 | **0.17** | **2.6** | <0.06 | <1.15 | <0.006 | <0.05 | <0.03 | <0.14 |
| **8** | 7,96 | 809 | 185,5 | <0.03 | <0.5 | <0.01 | <0.15 | <0.06 | <1.15 | <0.006 | <0.05 | <0.03 | <0.14 |

### Table S7. Comparison between Pb detection methods

| *Technique* | *Pro* | *Con* |
|---|---|---|
| | ***Atomic Quantification*** | |
| *Atomic Spectroscopy*[1] | LOD depends on ionization technique. (GFAAS: 1nM). Less expensive than ICP-MS<br>Detects Total dissolved element. | Requires sample pretreatment ( Sample Filtration, Acidification, Calibration). Matrix interferences. |
| *ICP-Mass Spectrometry*[1] | Lowest LOD (0.5 pM)<br>Detects Total dissolved element | Requires sample pretreatment Expensive facility and instrumentation. |
| | ***Electrochemistry*** | |
| *Voltammetry*[2,3] | EPA compatible LOD | Electrochemical specific equipment. Needs careful calibration. |
| *Portable Voltammetry*[2,3] | Portable | Especially sensitive to sample, reaction and equipment conditions |
| ***Spectroscopy*** | | |
| *Colorimetry*[4] | Inexpensive portable spectrophotometer | Not regulation compatible LOD (2 μM) |
| | ***"at home" tests kits*** | |
| *Lead paint test*[5] | EPA approved lead paint testing kits | Not compatible with water |
| *Water lead testing strips*[6] | - | Not EPA approved |
| | ***Synthetic Biology*** | |
| *Whole Cell Biosensors (WCBs)*[7] | Cell engineering enables signal modulation, crosstalk fixing and interference. Compatible with multiple signal outputs (colorimetric, fluorescence, luminescence) | LOD varies with circuit and signal output but typically is not compatible with current regulation. Biocontainment. Requires cell growth times and conditions. |
| *Cell Free Biosensors: ROSALIND*[8] | Low sample volume. Needs only inexpensive illuminator. Highly stable and portable. | Not regulation compatible LOD (1.25μM) |
| *Cell Free Biosensors: ROSALIND - ARCTA*[9] | Ultra-low LOD (6pM)* | Fluorescence levels are not shown to be compatible with an illuminator. |
| *Cell Free Biosensors: ROSALIND – NALF (this paper)* | Tunable LOD, matching EPA regulations (50nM). interpretable results. | Requires extra chromatography step. |

***** While the authors propose this LOD, the analytical performance of this type of sensor is not compatible with the quantitative analysis performed, particularly given that a linear response is not expected over such an extensive inducer range.

1. Bings, N. H., Bogaerts, A. & Broekaert, J. A. C. Atomic Spectroscopy: A Review. *Anal. Chem.* **82**, 4653–4681 (2010).
2. Gumpu, M. B., Sethuraman, S., Krishnan, U. M. & Rayappan, J. B. B. A review on detection of heavy metal ions in water – An electrochemical approach. *Sensors and Actuators B: Chemical* **213**, 515–533 (2015).
3. Sulthana, S. F. *et al.* Electrochemical Sensors for Heavy Metal Ion Detection in Aqueous Medium: A Systematic Review. *ACS Omega* **9**, 25493–25512 (2024).
4. Chen, Z. *et al.* Colorimetric detection of heavy metal ions with various chromogenic materials: Strategies and applications. *Journal of Hazardous Materials* **441**, 129889 (2023).
5. *Lead Test Kits - EPA*. https://www.epa.gov/lead/lead-test-kits.
6. Analytical Methods Approved for Drinking Water Compliance Monitoring of Inorganic Contaminants and Other Inorganic Constituents.
7. Kim, H. J., Jeong, H. & Lee, S. J. Synthetic biology for microbial heavy metal biosensors. *Anal Bioanal Chem* **410**, 1191–1203 (2018).
8. Jung, J. K. *et al.* Cell-free biosensors for rapid detection of water contaminants. *Nature Biotechnology* **38**, 1451–1459 (2020).
9. Zhang, X. *et al.* New Rolling Circle Transcription Based on Allosteric Transcription Factors (aTFs) for Trace Detection of Water Contaminants. *Anal. Chem.* acs.analchem.5c00217 (2025) doi:10.1021/acs.analchem.5c00217.

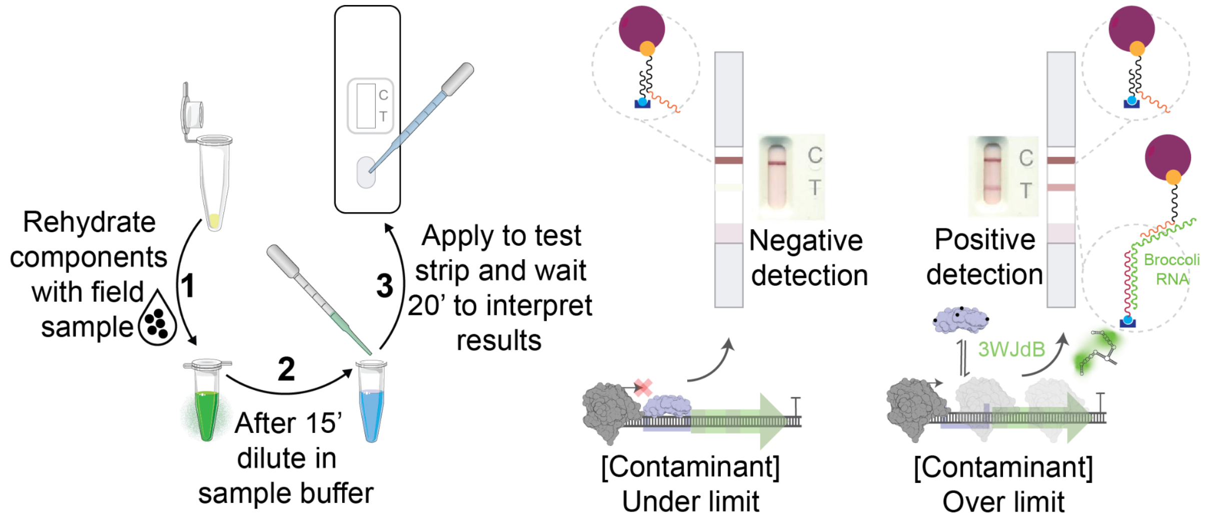

**Figure S1.** Schematic representation of the ROSALIND-NALF assay. The assay begins by rehydrating a freeze-dried reaction with 20 µL of sample, which is gently mixed until fully solubilized. After 20 minutes, a 1 to 5 µL aliquot of the reaction is taken and diluted with sample buffer and IVT buffer to a final volume of 100 µL. This 100 µL mixture is applied to a test strip, and results are interpreted after 15 minutes. A single control line indicates the contaminant concentration is below the test's detection limit, the presence of both a control and test line indicates a concentration above the limit, and a test line without a control line signifies an invalid test.

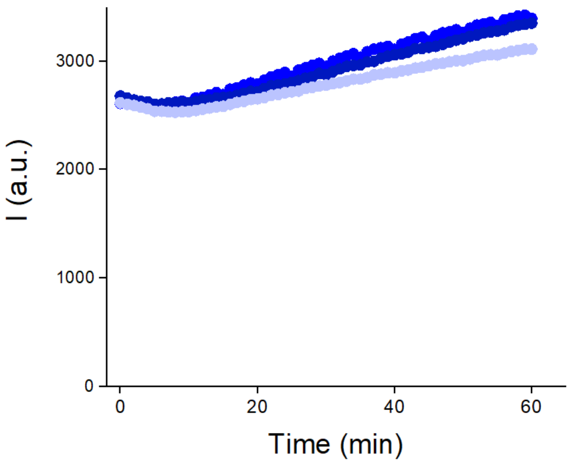

**Figure S2.** Disruption of an IVT reaction with kleptamers, without stopping transcription. Addition of 50 pmol of KbFull to 10 µL of an in vitro transcription reaction initially causes a decrease in fluorescence intensity, followed by a gradual increase as transcription shifts the equilibrium. Three biological replicates are shown as blue dots in varying shades.

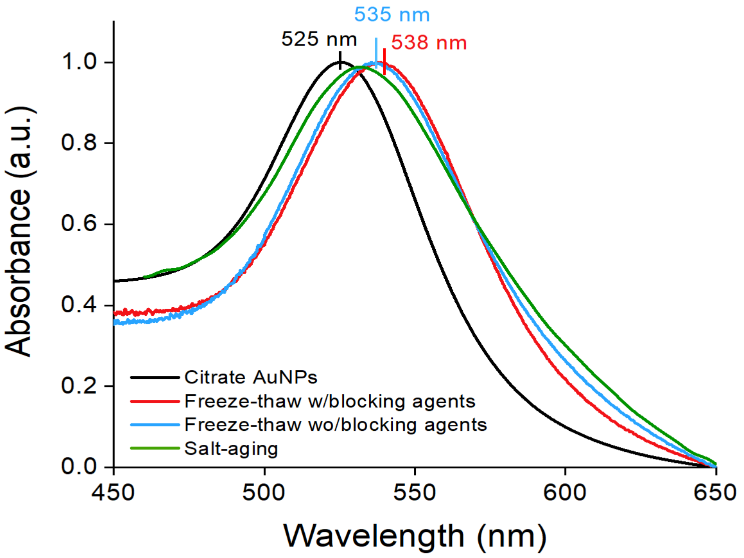

**Figure S3.** UV-vis spectra of citrate-capped AuNPs and DNA-AuNP conjugates synthesized via different methods. The black curve corresponds to citrate-capped 30 nm AuNPs, with a localized surface plasmon resonance (LSPR) peak at 525.4 nm. The red and blue curves represent DNA-AuNP conjugates prepared using the freeze-thaw method, with and without blocking agents, respectively. The green curve shows the conjugate synthesized using the salt-aging method. In all cases, a shift to longer wavelengths in the LSPR peak indicates surface modifications of the AuNPs. Absorbance values are normalized.

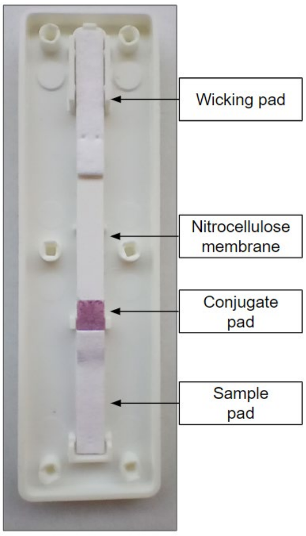

**Figure S4.** Representative strip showing the full assembly of all its components.

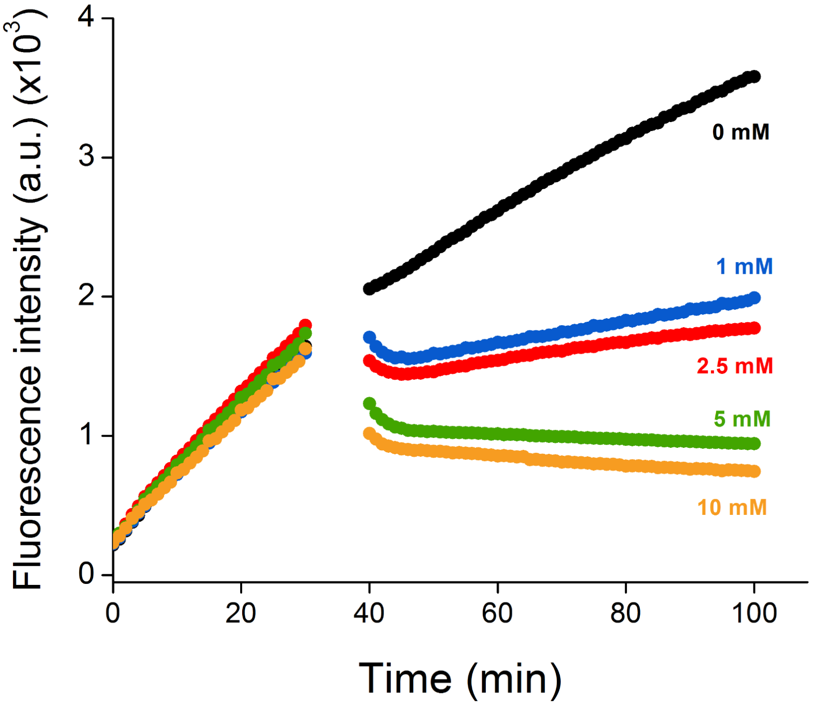

**Figure S5.** Effect of the addition of different concentrations of EDTA to the IVT reaction product. A decrease in fluorescence intensity is observed as the EDTA concentration increases.

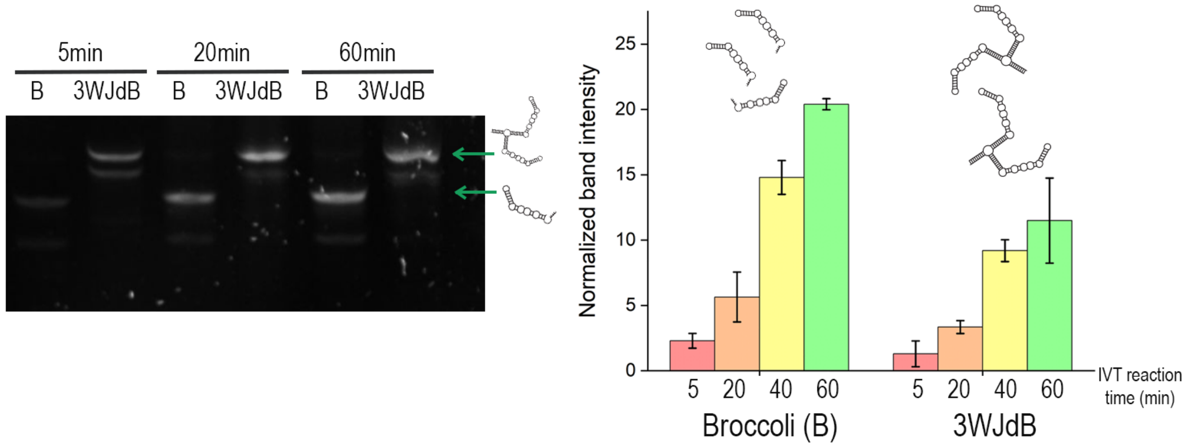

**Figure S6.** 10% dPAGE analysis of the IVT products at various reaction times for two different DNA templates: one containing a single Broccoli aptamer (B) and the other a 3WJ divalent Broccoli aptamer (3WJB). The normalized band intensity was calculated by adjusting for the number of nucleotides in each template, accounting for DNA length. Right: Normalized band intensities at different IVT reaction times (5, 20, 40, and 60 minutes) for both templates. The Broccoli template produced a higher amount of RNA compared to the 3WJ divalent Broccoli template (1.5X).

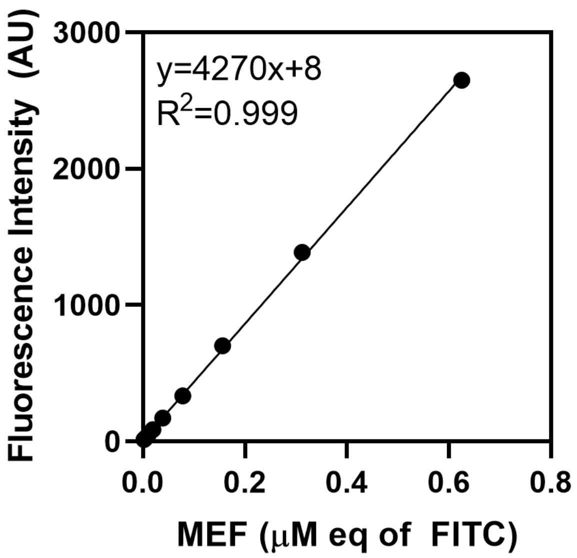

**Figure S7.** FITC calibration. Standard calibration curve used to obtain the conversion factor between μM equivalents of FITC and the fluorescence intensity measured with the corresponding experimental setup.

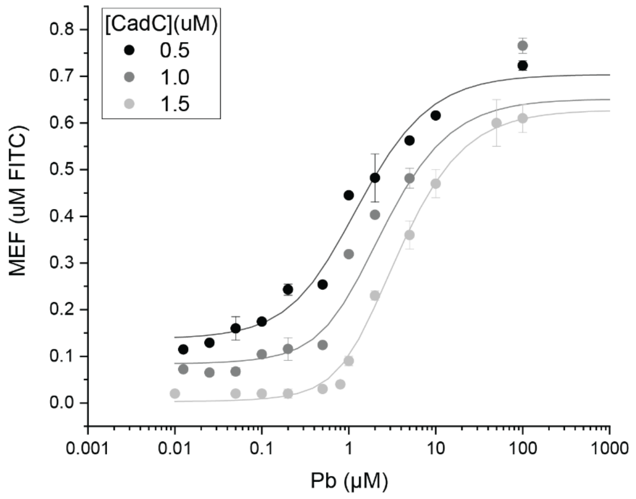

**Figure S8.** Standardized fluorescence (MEF) measurements for IVT reactions were performed with varying concentrations of Pb(II) and three different CadC concentrations at 25nM DNA template concentration. The lines represent the fit to the model depicted in figure 3A, increasing CadC concentration in these conditions also show some inhibition of the T7 activity, thus these datasets were not included in estimating the parameters in **Table S5**.

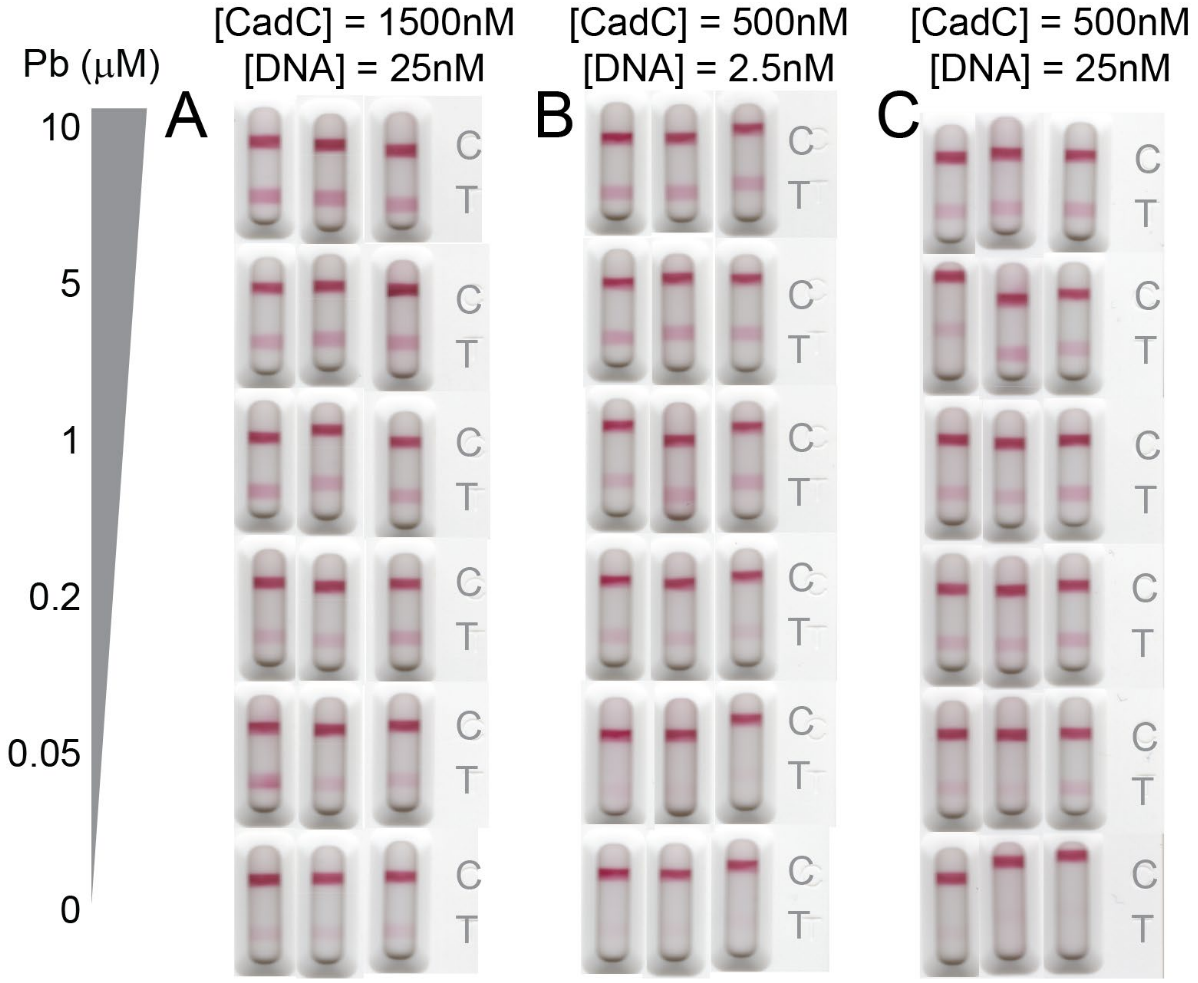

**Figure S9.** Replicates of the ROSALIND-NALF Pb(II) detection assays shown in Figure 3. Lateral flow strips display the colorimetric output from IVT reactions regulated by the CadC transcription factor under three different assay conditions: **A.** 1500 nM CadC with 25 nM DNA template, **B.** 500 nM CadC with 2.5 nM DNA template and **C.** 500 nM CadC with 25 nM DNA template. Reactions were performed with increasing concentrations of Pb(II) (0–10 μM, indicated on the left) and stopped with 5 mM EDTA, prior to dilution and application to the strips as described in Figure 3.

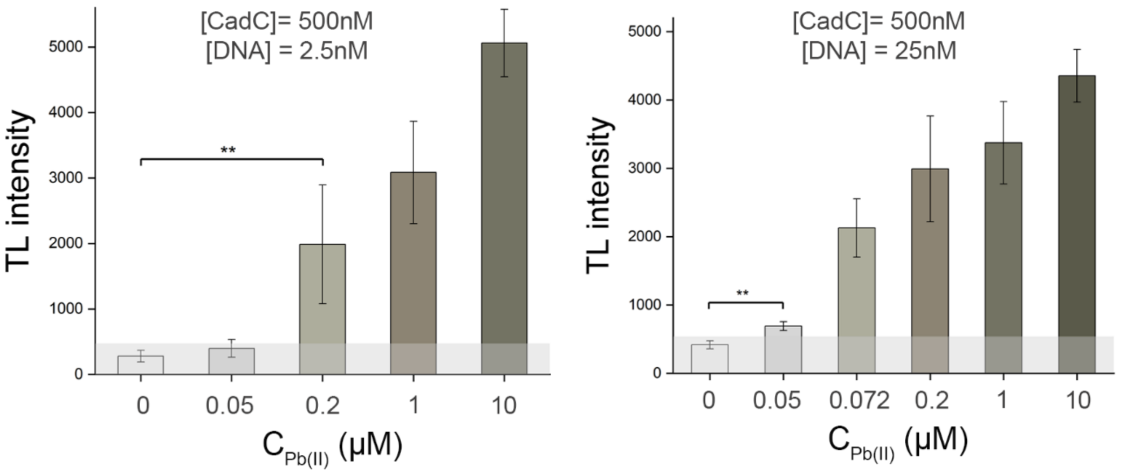

**Figure S10.** Detection limit determination. ROSALIND-NALF assay results for an optimized assay performed with 2.5 nM DNA template and 0.5 μM CadC (left) and ROSALIND-NALF assay performed with 25 nM DNA template and 0.5 μM CadC. two-tailed Student's t test; **p<0.05, bars represent mean ± SD.

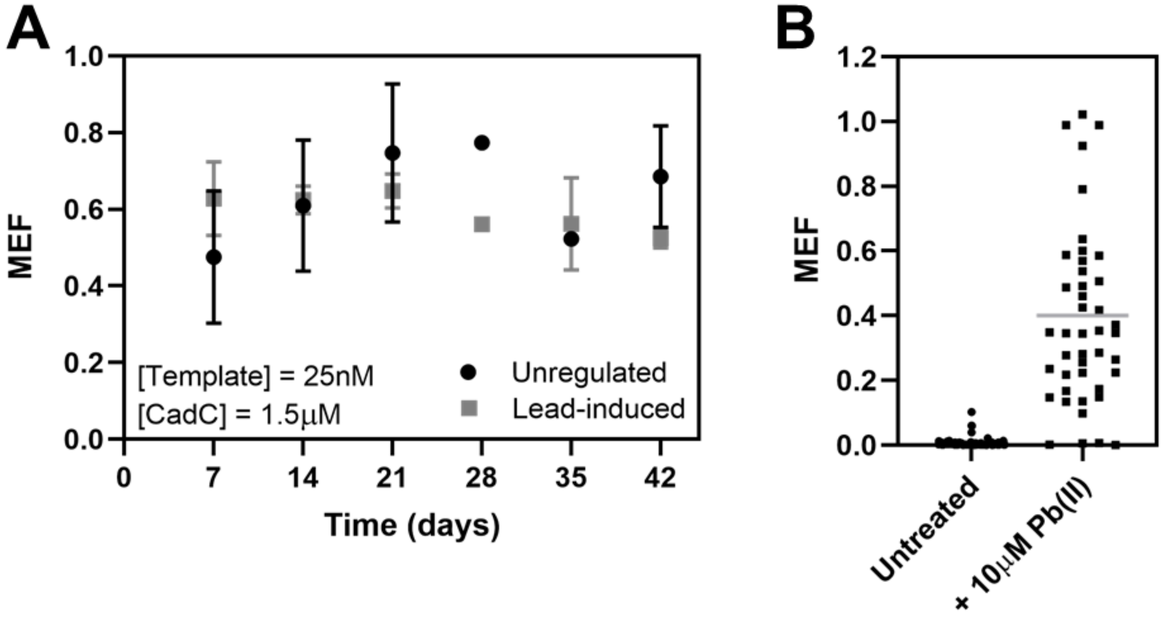

**Figure S11. A.** Stability of lyophilized stored ROSALIND reactions over time. ROSALIND reactions were lyophilized overnight, then packaged in vacuum-sealed bags with a desiccant and oxygen absorber, stored in dark containers until rehydration. With this conservation method, lyophilized reactions are functional for at least 1.5 months, showing stable signal over weeks. Though lead-induced reactions present a slight decay, the signal is clearly visible. Unregulated reactions were lyophilized with 25 nM of the template and lead-induced reactions with 25 nM of the template and 1.5 μM of the transcriptional repressor. Unregulated and lead-induced reactions were rehydrated with laboratory-grade water. **B.** Results from ROSALIND reactions (25nM template; 1.5µM CadC) done with untreated water from 53 monitoring sites across the Matanza Riachuelo Basin compared to 44 samples spiked with 10µM Pb(II). The rise in signal observed across the majority of the samples indicates that ROSALIND is compatible with testing superficial water samples although the spread among the intensities obtained reveals an important matrix effect.

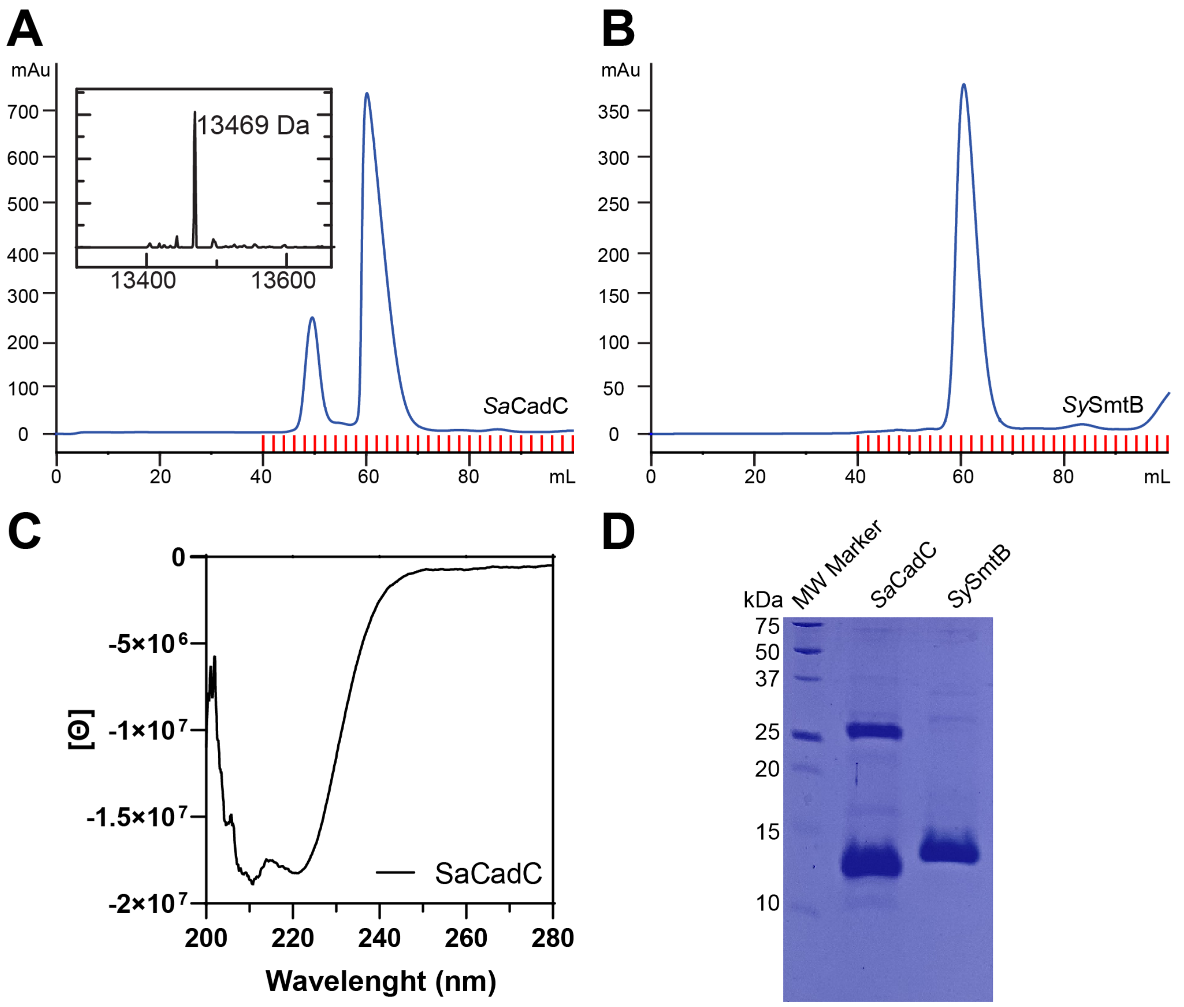

**Figure S12.** Transcription factor purification quality controls. **A.** and **B.** Chromatograms from size exclusion chromatography (Superdex S75) performed as the last purification step of *Sa*CadC and *Sy*SmtB. The inset in A. corresponds to an LC-ESI-TOF mass spectrometry measurement to corroborate the identity of the majoritarian SEC peak as *Sa*CadC. The mass of the intact protein corresponds to what is expected for the K2E mutant of the annotated sequence (UNIPROT id: P20047). **C.** Circular dichroism spectra of SaCadC corresponds to a correctly folded protein. **D.** SDS-PAGE (14%) of *Sa*CadC and *Sy*SmtB

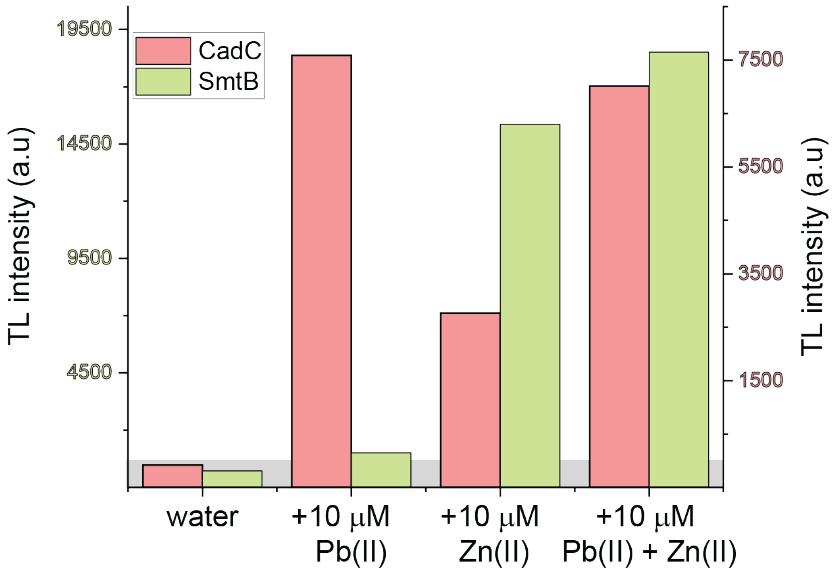

**Figure S13.** Comparison of test line intensities for CadC- and SmtB-based ROSALIND-NALF sensors in the presence of high concentration of divalent cations. The bar chart shows the LFA test line intensities for samples spiked with 10 µM Pb(II), 10 µM Zn(II), or a mixture of 10 µM Pb(II) + 10 µM Zn(II), evaluated using lateral flow readouts for both CadC and SmtB-based sensors. CadC, sensitive to both Pb(II) and Zn(II), shows detectable test line intensities for all conditions, with a stronger response to Pb(II). SmtB, which has a higher specificity for Zn(II), exhibits a strong response to Zn(II) and negligible interference from Pb(II). These results highlight the selectivity difference between the two sensors and suggest that SmtB can effectively distinguish high Zn(II) from high Pb(II) content.

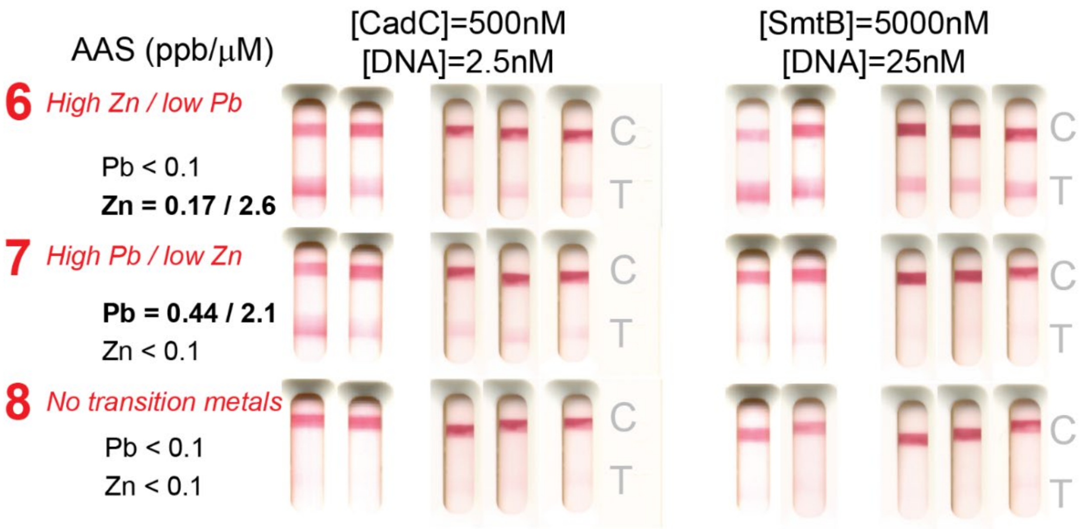

**Figure S14:** Replicates of the ROSALIND-NALF Pb(II) detection assays results shown in Figure 5D. Each set shows repeated tests of field water samples with the indicated Pb(II) and Zn(II) concentrations (determined by AAS) using CadC-based (*left*) and SmtB-based (*right*) ROSALIND-NALF sensors.

Standard Methods, n.d. Standard Methods: 3030 E:  Nitric Acid Digestion of Metals https://doi.org/10.2105/SMWW.2882.041

Standard Methods, n.d. Standard Methods: 3111B:  Metals in water by FLAA. https://doi.org/10.2105/SMWW.2882.043

Standard Methods, n.d. Standard Methods: 2340C:  EDTA Titrimetric Method (Hardness) https://doi.org/10.2105/SMWW.2882.025.

Thavarajah, W., Owuor, P.M., Awuor, D.R., Kiprotich, K., Aggarwal, R., Lucks, J.B., Young, S.L., 2023. The accuracy and usability of point-of-use fluoride biosensors in rural Kenya. npj Clean Water 6, 5. https://doi.org/10.1038/s41545-023-00221-5

Thavarajah, W., Verosloff, M.S., Jung, J.K., Alam, K.K., Miller, J.D., Jewett, M.C., Young, S.L., Lucks, J.B., 2020. A primer on emerging field-deployable synthetic biology tools for global water quality monitoring. npj Clean Water 3, 18. https://doi.org/10.1038/s41545-020-0064-8

Tripodi, M.A., Cueto, G.R., Muschetto, E., Hancke, D., Suárez, O.V., 2023. Intra- and inter-annual variations in metal concentrations in the superficial water of a highly polluted urban basin of Argentina. Environ Sci Pollut Res 30, 60838–60853. https://doi.org/10.1007/s11356-023-26391-w

Turner, J., 1996. Zn2+-sensing by the cyanobacterial metallothionein repressor SmtB: different motifs mediate metal-induced protein-DNA dissociation. Nucleic Acids Research 24, 3714–3721. https://doi.org/10.1093/nar/24.19.3714

WHO (World Health Organization), n.d. Water, sanitation and hygiene: burden of disease. https://www.who.int/data/gho/data/themes/topics/water-sanitation-and-hygiene-burden-of-disease (accessed May 2025)

WHO/UNICEF Joint Monitoring Programme for Water Supply, Sanitation and Hygiene (JMP), 2021. Progress on household drinking water, sanitation and hygiene 2000–2020: Five years into the SDGs.

Yang, Y., Li, W., Liu, J., 2021. Review of recent progress on DNA-based biosensors for Pb2+ detection. Analytica Chimica Acta 1147, 124–143. https://doi.org/10.1016/j.aca.2020.12.056

Yetisen, A.K., Akram, M.S., Lowe, C.R., 2013. Paper-based microfluidic point-of-care diagnostic devices. Lab Chip 13, 2210. https://doi.org/10.1039/c3lc50169h
